# Lymphotoxin-alpha expression in the meninges causes lymphoid tissue formation and neurodegeneration

**DOI:** 10.1101/2021.04.27.441396

**Authors:** Rachel E James Bates, Eleanor Browne, Renee Schalks, Heather Jacobs, Li Tan, Puja Parekh, Roberta Magliozzi, Massimiliano Calabrese, Nicholas D. Mazarakis, Richard Reynolds

## Abstract

Lymphotoxin alpha (LTα) plays an important role in lymphoid organ development and cellular cytotoxicity in the immune system. LTα expression is increased in the cerebrospinal fluid of naïve and progressive multiple sclerosis (MS) patients and post-mortem meningeal tissue. Here we show that persistently increased levels of LTα in the cerebral meninges can give rise to lymphoid-like structures and underlying MS-like cortical pathology. Stereotaxic injections of recombinant LTα into the rat meninges leads to acute meningeal inflammation and subpial demyelination that resolves after 28 days. Injection of an LTα lentiviral vector induces lymphoid-like immune cell aggregates, maintained over 3 months, including T-cell rich zones containing podoplanin+ fibroblastic reticular stromal cells and B-cell rich zones with a network of follicular dendritic cells, together with expression of lymphoid chemokines and their receptors. Extensive microglial activation, subpial demyelination and marked neuronal loss occurs in the underlying cortical parenchyma. These results show that chronic LTα overexpression is sufficient to induce formation of meningeal lymphoid-like structures and subsequent neurodegeneration.

**Summary:** Increased release of lymphotoxin-alpha contributes to the pro-inflammatory milieu of the cerebrospinal fluid of MS patients. A persistent elevated expression of this cytokine in the meninges of rats gives rise to chronic inflammation with lymphoid tissue induction and accompanying neurodegenerative and demyelinating pathology in the underlying brain tissue.

## Introduction

The progressive stages of multiple sclerosis (MS) are characterised by an increasing burden of grey matter pathology that is suggested to play a major role in the irreversible accumulation of sensory, cognitive and motor symptoms (Calabrese et al., 2015a; Fisniku et al., 2008; Harrison et al., 2015; Kutzelnigg et al., 2005; Reynolds et al., 2011). Accumulating axon and neuronal loss that are stimulated by an ongoing compartmentalised inflammation are now widely accepted to be one of the main pathological correlates of clinical progression (Calabrese et al., 2015b; Magliozzi et al., 2010; Stadelmann, 2011). Recent studies on human post-mortem tissues have suggested that chronic inflammation in the leptomeninges may be one of the main drivers of cortical pathology (Haider et al., 2016; Howell et al., 2011; Magliozzi et al., 2007; Magliozzi et al., 2018). The presence of leptomeningeal inflammation and the development of tertiary lymphoid-like structures (TLS) in the subarachnoid space (Magliozzi et al., 2007; Serafini et al., 2004) correlate with a gradient of neuronal loss in the cortex that is associated with a shorter time to milestones of clinical progression, including a shorter time to wheelchair use, shorter disease duration and younger age at death (Howell et al., 2011; Magliozzi et al., 2010). These TLS are discrete aggregates of B cells and T cells, plasma cells, antigen-presenting cells and a network of follicular dendritic cells, display varying degrees of organisation and may serve as sites for autoantigen presentation and intrathecal antibody production, both of which are important in driving inflammation in chronic MS (Bell et al., 2019; Magliozzi et al., 2007; Serafini et al., 2004). Meningeal inflammation is seen from the very early stages of MS (Bevan et al., 2018; Lucchinetti et al., 2011) and in multiple CNS locations (Howell et al., 2015; Magliozzi et al., 2010; Reali et al., 2020), but the molecular and cellular mechanisms involved in the formation and maintenance of these immune cell aggregates remains unclear.

The presence of compartmentalised inflammation in the meninges and the underlying gradient of pathology in the cortical GM has given rise to the hypothesis that proinflammatory cytokines and other cytotoxic molecules in the CSF may diffuse into the underlying cortex and directly or indirectly initiate cellular damage (Calabrese et al., 2015b; Dendrou et al., 2015; Gardner et al., 2013; Magliozzi et al., 2007). This also raises the possibility that the same molecule(s) that are involved in stimulating the development of lymphoid tissues could also be responsible for the cortical cell damage. Analysis of the composition of post-mortem and patient CSF of MS cases with increased cortical pathology (Magliozzi et al., 2018; Magliozzi et al., 2020) has revealed the elevated presence of proinflammatory cytokines (eg. IFNγ, TNF, IL2, IL6), molecules related to B-cell activation (eg. BAFF, APRIL, LIGHT, IL10) and lymphoid neogenesis (eg. LTα, CXCL13, CXCL10). Of these molecules, lymphotoxin-α (LTα) is known to have a role in cellular cytotoxicity, lymphoid neogenesis and TLS formation (Gommerman and Browning, 2003; Gommerman et al., 2003; Kratz et al., 1996), but its involvement in the pathogenesis of MS has been little studied. Lymphotoxin-α can form a soluble homotrimer that binds to TNF receptors 1 and 2 (TNFR1/TNFR2) and it can also form heterotrimers together with LTβ, as either LTα2β1 or LTα1β2, which remain tethered in the cell membrane and bind to the LTβ receptor (LTβR). The involvement of LTα in the pathogenesis of MS was implicated by its ability to potently induce apoptosis of oligodendrocytes *in vitro* (Selmaj et al., 1991b), and its expression by CD3+ lymphocytes and microglia at the lesion edge of acute and chronic active MS lesions (Cannella et al., 1997; Raine et al., 1998; Selmaj et al., 1991a). Increased LTα mRNA expression was observed in peripheral blood lymphocytes prior to relapse in RRMS, and expression was strongly upregulated by stimulation with MBP *in vitro*, suggesting that LTα may play a role in the exacerbation of MS (Matusevicius et al., 1996; Rieckmann et al., 1995). Increased LTα expression has also been demonstrated in peripheral blood B-cells (Romme Christensen et al., 2013) and increased release by CD8+ T-cells in SPMS patients (Buckle et al., 2003).

LTα is known to play a key role in lymphoid tissue organogenesis and maintenance (Gommerman and Browning, 2003). Mice that are deficient in LTα have significant defects in the development of lymphoid organs, specifically lacking secondary lymphoid organs including lymph nodes (LN) and Payer’s patches (PP)(De Togni et al., 1994) and are resistant to experimental allergic encephalomyelitis induced in C57Bl6 mice by immunisation with MOG peptide 35-55 (Suen et al., 1997). In contrast, ectopic transgenic expression driven by the rat insulin promotor caused chronic inflammation and TLS formation in the kidney and pancreas (Kratz et al., 1996). Whilst LTα has also been shown to be a key player in formation of TLS in the lung during influenza and in synovial tissue in rheumatoid arthritis (Hirose et al., 2018), the role of LTα in chronic leptomeningeal inflammation and CNS TLS formation has yet to be examined.

Meningeal inflammation accompanied by various stages of TLS formation has been reported in several animal models of MS, including the relapsing remitting model of experimental autoimmune encephalomyelitis (rrEAE) in the SJL/J mouse immunised with PLP peptide (Magliozzi et al., 2004), chronic progressive EAE in the Biozzi-ABH mouse (Magliozzi et al., 2004), in C57Bl/6 mice immunised with an MBP-PLP fusion protein (Kuerten et al., 2012) and following adoptive transfer of MOG-specific Th-17 T-cells into C57Bl/6 mice (Mitsdoerffer and Peters, 2016; Peters et al., 2011; Pikor et al., 2015). However, little is known concerning the molecular mediators responsible for initiation and maintenance of such organised immune cell infiltrates and their role in the development of demyelinating and neurodegenerative pathology in the underlying CNS tissues. Our previous studies have demonstrated a close association between the formation of TLS in the subarachnoid space of the cerebral cortex and demyelinating and neurodegenerative pathology in the underlying grey matter, which is not reproduced by any of the EAE models (Howell et al., 2011; Magliozzi et al., 2010; Magliozzi et al., 2019). In order to study this relationship, we have recently developed a novel model in the rat in which the levels of pro-inflammatory cytokines can be chronically and ectopically raised to pathological levels (James et al., 2020). Given the potential role of LTα in both cellular cytotoxicity and TLS formation, here we aimed to show whether acute or chronically elevated levels of LTα in the subarachnoid space of the rat brain can result in meningeal TLS formation and subsequent demyelination and neuronal loss. Establishing the factors that may drive the formation of TLS in MS and their causal relationship to cortical pathology will open up an avenue for the development of an effective approach to slowing the progressive course of MS.

## Results

### Expression of LTα in MS CSF and meninges

A cohort of recently diagnosed drug naïve MS patients were grouped based on the presence of at least 10 cortical lesions (MS-high) or less than 2 lesions (MS-low) from 3D DIR-MRI imaging (cohort described in Magliozzi et al., 2018). Significantly increased levels of LTα were present in the CSF of the MS-high group compared to controls and the MS-low group (Fig. 1A). In a similar way, significantly elevated LTα protein levels (Fig 1B) were found in the CSF of the MS-high post-mortem SPMS brains (>60% GM demyelination and high levels of immune cell infiltration; Magliozzi et al., 2018), compared to both controls and the MS-low group (<40% GM demyelination and a low density of diffuse meningeal inflammation). Levels of LTα mRNA in the meninges were also significantly increased in SPMS brains compared to controls (3.2 ± 1.2 fold; p=0.028) (Fig. 1C). Immunostaining for LTα showed cytoplasmic staining in cells within the meningeal infiltrates (Fig. 1D), many of which co-stained for the pan T-cell marker CD3 (Fig. 1E-G). Immunohistochemistry for the LTβR showed a diffuse pattern of reticular like staining throughout the meningeal infiltrates (Fig. 1I).

**Figure 1.**
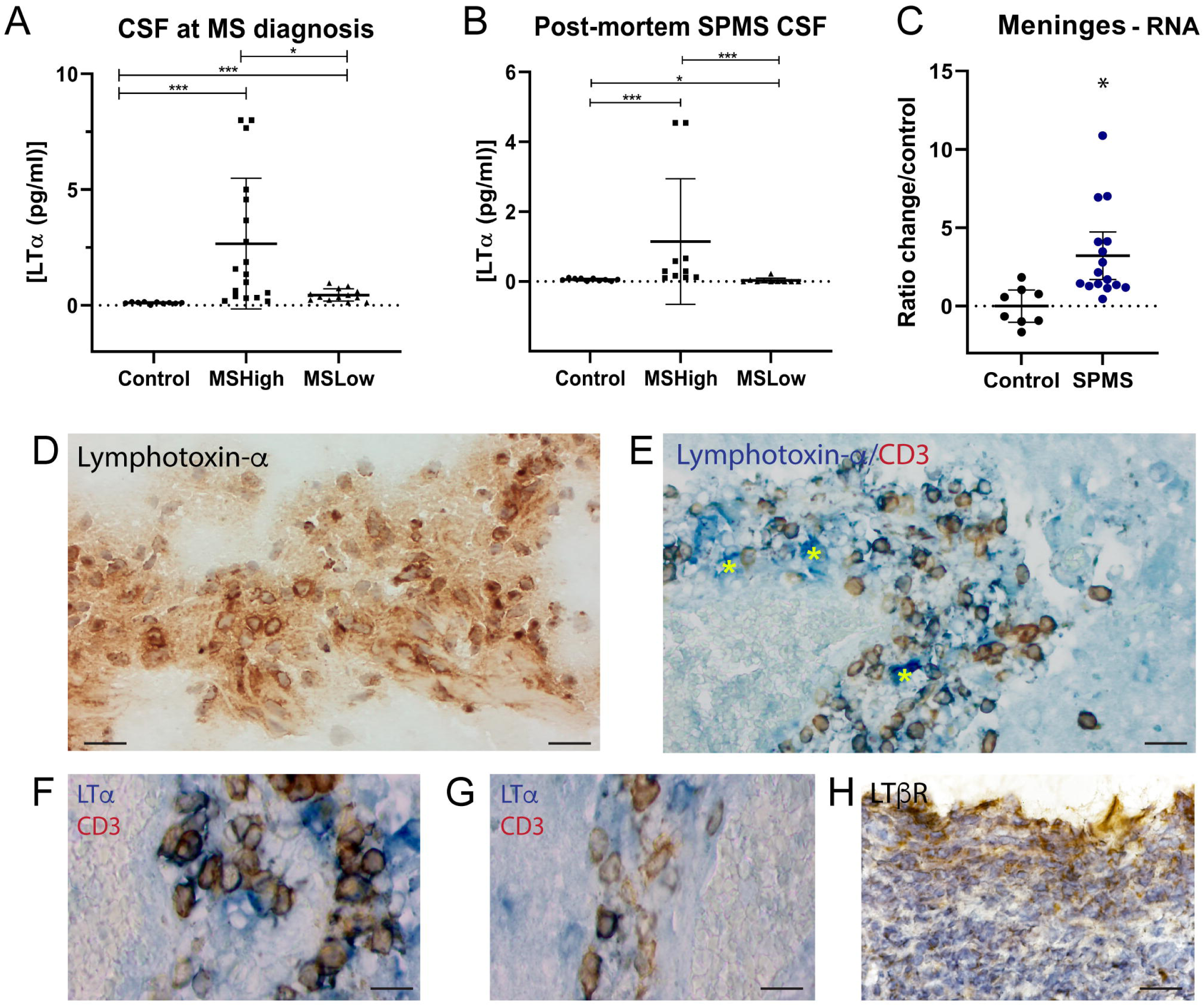
Analysis of the levels of LTα gene transcript and protein. **(A)** Levels of LTα measured in diagnostic CSF samples from 10 non-inflammatory neurological controls, 18 MSHigh and 13 MSLow cases by Bioplex analysis. The number of cortical lesions on 3D DIR MRI was used to stratify naïve MS patients as MSHigh (greater than 10 cortical lesions) or MSlow inflammation (less than 2 cortical lesions). **(B)** LTα protein levels in post-mortem CSF from 10 MSHigh, 10 MSLow and 10 control cases, measured by electrochemoluminescence (MSD). SPMS post-mortem cases were classed as MSHigh or MSLow based on the degree of immune cell infiltration in the meninges, based on the numbers of haematoxylin nuclei in the infiltrate. **(C)** LTα gene transcript levels in meninges dissected from post-mortem tissue blocks of 16 SPMS cases and 8 controls, determined by QPCR. Graph shows fold change in mRNA levels in MS versus control. **(D)** Immunostaining for LTα in meningeal aggregates at 10x shows cytoplasmic staining in numerous cells in meningeal infiltrates. Scale bar = 50µm. **(E)** Co-staining for lymphotoxin-α (blue) and CD3 (brown) in meninges show double positive staining on CD3+ T-cells. In addition, some LTα expressing cells are not CD3+ T-cells (yellow stars). Scale bar = 50µm. **(F-G)** LTα (blue) and CD3 (brown) co-staining at higher magnification. Scale bar=20µm. **(H)** Immunostaining for LTβR in meningeal aggregates shows a reticular like staining. Statistics: 1-way analysis of variance with Tukey post-test.

### Meningeal inflammation after recombinant cytokine injection in CSF

The formation of dense cellular infiltrates, down the entire depth of the sulcus and extending across the lateral surface of the cortex was observed between 3-14 days following cytokine injection, with the greatest levels seen between 3-7 days, but were absent in controls that received only a PBS injection (Fig. 2A). Very few immune cells were seen in the underlying cortical parenchyma at any timepoint. The meningeal infiltrate was composed of predominantly CD4+ and CD8+ T cells and CD79a+ B-cells (shown in Fig. 2B at 3dpi). CD4+ and CD8+ T-cells were highest at 3dpi and decreased thereafter until they were almost absent by 14 days (Fig. 2C-D). At both 3 and 7dpi the numbers of CD4+ cells were approximately twice the numbers of CD8+ T-cells (Fig. 2C-D). Infiltration of CD79a+ B-cells was maximal at 7 days post-injection with numbers staying significantly high at 14 and 21 dpi (Fig. 2E). Whilst CD4+ and CD8+ T-cells were roughly evenly distributed throughout the space with no observable organisation, CD79α+ B-cells often formed discrete, dense clusters of cells (Fig. 2F). Approximately 65% of CD79a+ B-cells co-expressed the proliferation marker Ki67 at 3dpi with the percentage of CD79a+Ki67+ cells decreasing over time (Fig. 2G-H).

**Figure 2.**
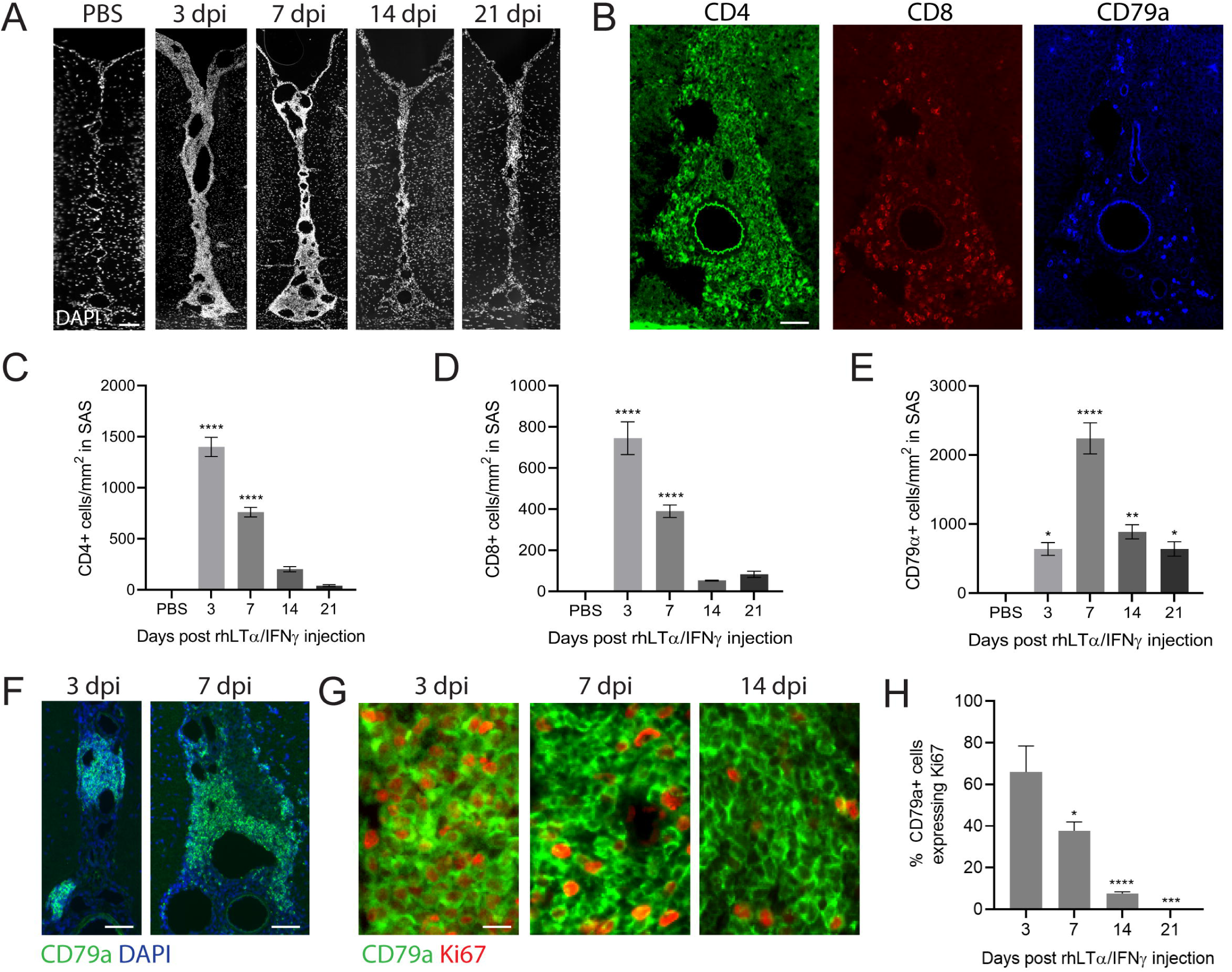
Injection of recombinant LTα and IFNγ into the subarachnoid space of rats leads to meningeal inflammation. **(A)** Cytokine injection into the SS led to the formation of large dense infiltrates of cells the length of the sagittal sulcus and across the surface of the cortex visualised here at 4x magnification by DAPI immunostaining. Injection of PBS did not lead to any infiltration. Scale bar = 100um. **(B)** Representative images of CD4, CD8 and CD79a from sequential sections at 3dpi post injection of cytokines. Scale bar 50=μm. **(C-D)** Quantification of CD4+ (C) and CD8+ (D) cell numbers in the SS expressed as cells/mm^2^. Both CD4+ and CD8+ numbers were maximal at 3dpi with CD4+ cell number 104% higher than CD8+. 14 and 21dpi were not significantly increased compared to naïve. Data presented as mean ± SEM (n=3-4 per group). **** P<0.0001 cytokine compared to naïve. **(E)** Quantification of CD79a cells expressed as cells/mm^2^ in the sagittal sulcus. The maximal infiltration of CD79a+ B-cells into the sagittal sulcus was seen at 7dpi and was significantly highly than naïve at 3, 14 and 21dpi. * p<0.05; ** p<0.001, **** p<0.0001 cytokine vs naïve. **(F)** CD79a+ B-cells formed densely packed, focal clusters of cells within the meningeal aggregates. Scale bar=100µm. **(G)** Following cytokine injection a proportion of CD79a+ cells (green) were positive for the proliferation marker Ki67 (red). **(H)** Data showing the percentage of CD79a+ cells that co-stained with Ki67. At 3dpi 65% of cells were double positive (n=4), significantly decreasing to 37.7% at 7dpi (n=4), 7.5% at 14dpi (n=3) and 0.07% at 21dpi (n=3). Statistics: 1-way analysis of variance with Tukey post-test. *p<0.05, ***p<0.001, ****p<0.0001 compared to 3dpi.

### Demyelination and microglial activation after acute cytokine injection

Injection of PBS into the SS of control animals, either naive or those immunised with MOG+IFA or IFA only, did not result in any demyelination or activation of microglia (Fig. 3A-C). Rats immunised with low dose rmMOG in IFA that also received a stereotaxic injection of 1μg LTα and 75ng IFNγ recombinant proteins into the SS exhibited subpial demyelination of the upper cortical layers and midline layers as early as 3 dpi (Fig. 3D-F), which became even more extensive at 7dpi (Fig. 3G-I) and 14dpi (Fig. 3J-L) and was accompanied by substantial IBA1+ microglial activation. Demyelination extended from the corpus callosum boundary of the sagittal sulcus to the dorsal surface of the cortex and laterally from the midline across the subpial surface of the cortex through to rostral regions. By 21dpi there was very little apparent loss of MOG staining with only small patches of loss seen mostly in midline regions (Fig. 3M), with negligible IBA1+ microglial activation (Fig 3N-O), suggesting that remyelination had occurred. Quantification of the loss of MOG staining demonstrated that demyelination was greatest at 7dpi when compared to naïve animals with less demyelination at 3 and 14dpi in midline layer I (Fig. 3P) and midline layer II-III (Fig. 3Q). A similar pattern was seen in cortical layers I-IV (Fig. 3R) with significantly less MOG+ myelin staining at 7dpi (15.2 ± 2.25%) than at 3dpi (23.2 ± 3.9%) and 14dpi (25 ± 3%). By 21dpi there was no quantifiable demyelination in the cortical layers (Fig. 3R), but still some loss of MOG staining in midline regions (Fig. 3P-Q). Quantification of IBA1+ immunofluorescence showed levels peaked at 7dpi in cortical (Fig. 3S) and midline (Fig. 3T) layers, corresponding to the peak in demyelination, but were not significantly different from PBS animals by 21dpi.

**Figure 3.**
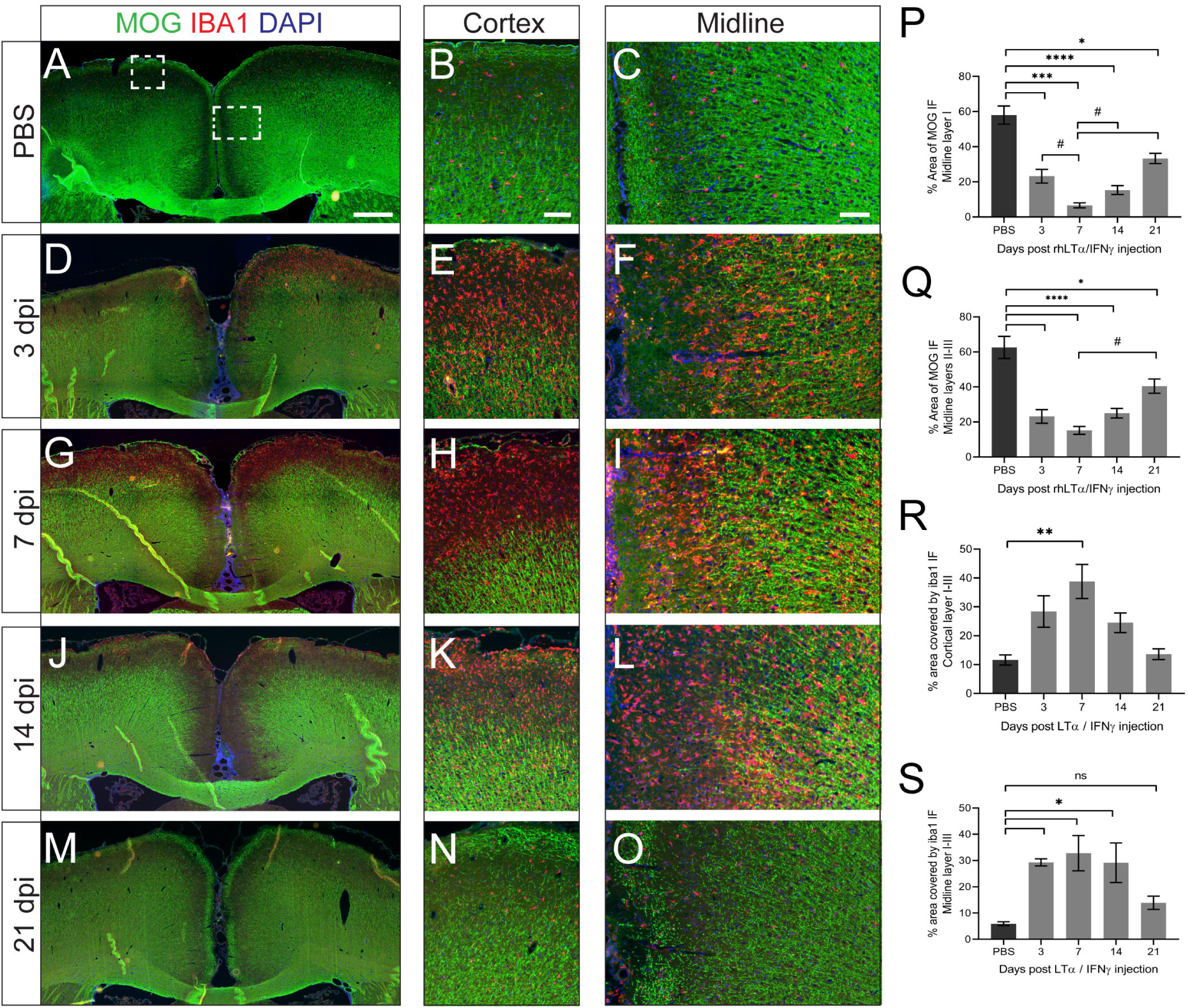
Subpial demyelination following injection of recombinant LTα+IFNγ into the subarachnoid space. **(A-O)** Immunofluorescence for myelin oligodendrocyte glycoprotein (MOG) and the microglial marker IBA1 in rmMOG immunised rats injected with PBS **(A-C)** or 1mg LTα and 75ng IFNγ recombinant cytokines culled at 3 **(D-F)**, 7 **(G-I)**, 14 **(J-L)** and 21 **(M-O)** days post-cytokine injection (dpi), demonstrating the degree of demyelination and microglial activation at various times points. Cortex and midline images are magnifications of the regions shown by the boxed areas in image A. Scale bars: A = 500µm, B and C = 50µm. **(P-Q)** Quantification of subpial demyelination at the injection site in layer I (P) and layers II-III **(Q)**, presented as the percentage area covered by MOG immunofluorescence (IF) (n=3-4 each group). Data is presented as the mean ± SEM. Statistics: 1-way analysis of variance with Tukey post-test. * P<0.01; *** p<0.005, **** p<0.001 cytokine vs PBS; # p<0.05 7 vs 21 dpi. (R-S) Measurement of the area covered by IBA1 immunofluorescence in cortical layers I-III **(R)** and midline layers I-III **(S)** and presented at mean ± SEM. * p<0.05; ** p<0.001 cytokine vs naïve.

#### Expression of the LT*α* transgene

In order to study the chronic effects of increased LTα expression by cells in the meningeal space, we developed a high titre viral vector. Primary meningeal cells could be 100% transduced with the lentiviral vectors for LTα and GFP, demonstrating a good tropism for the virus (Supplementary Fig 1A-F) with no obvious toxicity (Supplementary Fig. 1G). Transduced meningeal cells continued to produce transgene at increasing levels for several days (Supplementary Fig. 1K-L). Following stereotaxic injection into the subarachnoid space of the sagittal sulcus, as indicated by the presence of monastral blue dye at the injection site, mRNA for the LTα transgene was highly expressed at 90 days after injection in meninges dissected from near the injection site (Fig. 4A). Meningeal cells lining the sagittal sulcus were successfully transduced, as shown by the expression of GFP (Fig. 4B). Strong GFP expression could also be detected in meningeal cells lining the outer surface of the cortex. Immunofluorescence with anti-human LTα antibody demonstrated high levels of expression of the human LTα transgene in transduced meningeal cells down the sagittal sulcus at 28 and 90dpi (fig. 4C). Protein expression from the LTα transgene was detectable in CSF samples and maintained at 28 and 90 days whilst levels were undetectable in GFP vector injected animals (Fig. 4D). Human LTα protein was expressed at high levels in tissue taken from around the injection site, it was also present at low levels in surrounding cortical, but not cerebellum, tissue suggesting some transduction in the cortex, possibly via diffusion down perivascular spaces (Fig. 4E).

**Figure 4.**
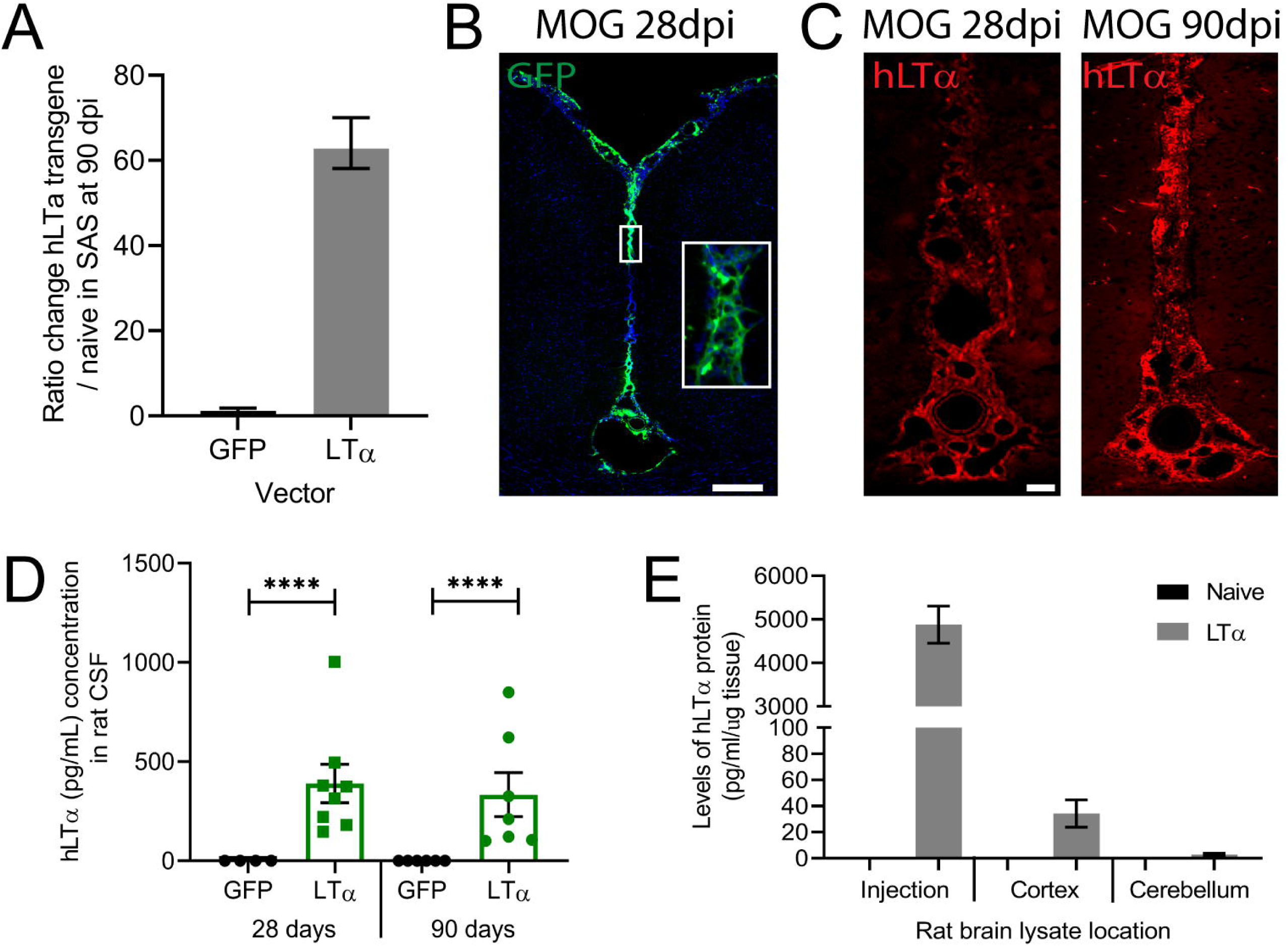
Primary meningeal cells transduced with lentiviral vector expressing human LTα. **(A)** Lentiviral vectors expressing eGFP as a control or LTα were injected into the sagittal sulcus. Following LV-LTα injection the human LTα transgene could be measured in the meninges dissected from rats at 90 dpi, demonstrating long term expression of the gene by the CMV promoter within the transduced meningeal cells (n=3 rats), but no expression in the eGFP controls. Data shown as ratio change of 2^ΔΔCT for hLTα mRNA in LVGFP and LVLTα compared to naïve animals. **(B)** Significant transduction of meningeal cells with the lentiviral vectors could be demonstrated by expression of the GFP protein (green), the length of the SS and across the lateral surface of the cortex. Higher magnification image in box. **(C)** Immunostaining for hLTα (red) showed expression of the human LTα protein along the length of the SS that was present both at 28 and 90 dpi demonstrating widespread transduction. **(D)** CSF samples were taken from the cisterna magna of injected animals shortly before perfusion at 28 and 90 dpi. hLTα levels were measured in 100μl CSF samples using electrochemiluminescence (MSD plates). hLTα could be detected at both 28dpi (mean 389.6pg/mL) and 90dpi (333.6pg/mL) at similar levels but was not present in LVGFP animals. Statistics: 1-way analysis of variance with Tukey post-test. **** p<0.001. **(D)** Quantification of hLTα protein levels by ELISA in tissue samples dissected from either naïve or LVLTα injected animals at 90 dpi at the site of injection and from the surrounding cerebral cortex and the cerebellum, demonstrating very local transduction.

### Meningeal inflammation after LTα overexpression in the subarachnoid space

In LTα lentiviral vector injected animals, substantial meningeal infiltrates developed and completely filled the midline subarachnoid space by 28 dpi and were maintained until at least 90 dpi (Fig. 5A), the latest time point studied. No qualitative difference was seen between IFA only and IFA+MOG immunised animals. CD4+ and CD8+ T-cells were roughly evenly distributed throughout the space, whereas CD79a+ B-cells tended to cluster more (Fig. 5B). IBA1+ myeloid cells were also present throughout the aggregates. Quantification of CD4+ and CD8+ T-cell numbers showed no significant difference in numbers between IFA and IFA+MOG animals at both 28 and 90 dpi (Fig. 5C-D), with approximately equal numbers of CD8+ and CD4+ T-cells. Although numbers of T-cells were lower at 90dpi, this still represented substantial infiltration. There was no difference in the number of CD79a+ B-cells between IFA or IFA+MOG animals at 28 or 90dpi (Fig. 5E) and numbers were similar to both CD4+ and CD8+ T-cells. Similar to the animals injected with recombinant cytokines, after viral vector injection CD79a+ B-cells formed into densely packed focal clusters (Fig. 5F), with immunoglobulin-expressing plasma cells sparsely distributed in the same areas (Fig. 5G). Few or no CD8+ or CD4 T-cells were present within these CD79a+ clusters and a proportion of CD79a+ cells were proliferating at both time points, as demonstrated by co-expression with Ki67 (Fig. 5H). The percentage of Ki67+ B-cells was stable between 28 and 90dpi and was not different between MOG+IFA and IFA only animals, suggesting a consistent level of B-cell proliferation that was not driven by the anti-MOG immune response (Fig. 5I-J). Large aggregates of infiltrating cells were also found along the lateral surface of the cortex (Fig. 5K), composed of CD4+ and CD8+ T-cells (Fig. 5L) and CD79a+ B-cells (Fig. 5M), and were present as T-cell rich regions and B-cell clusters (Fig. 5N).

**Figure 5.**
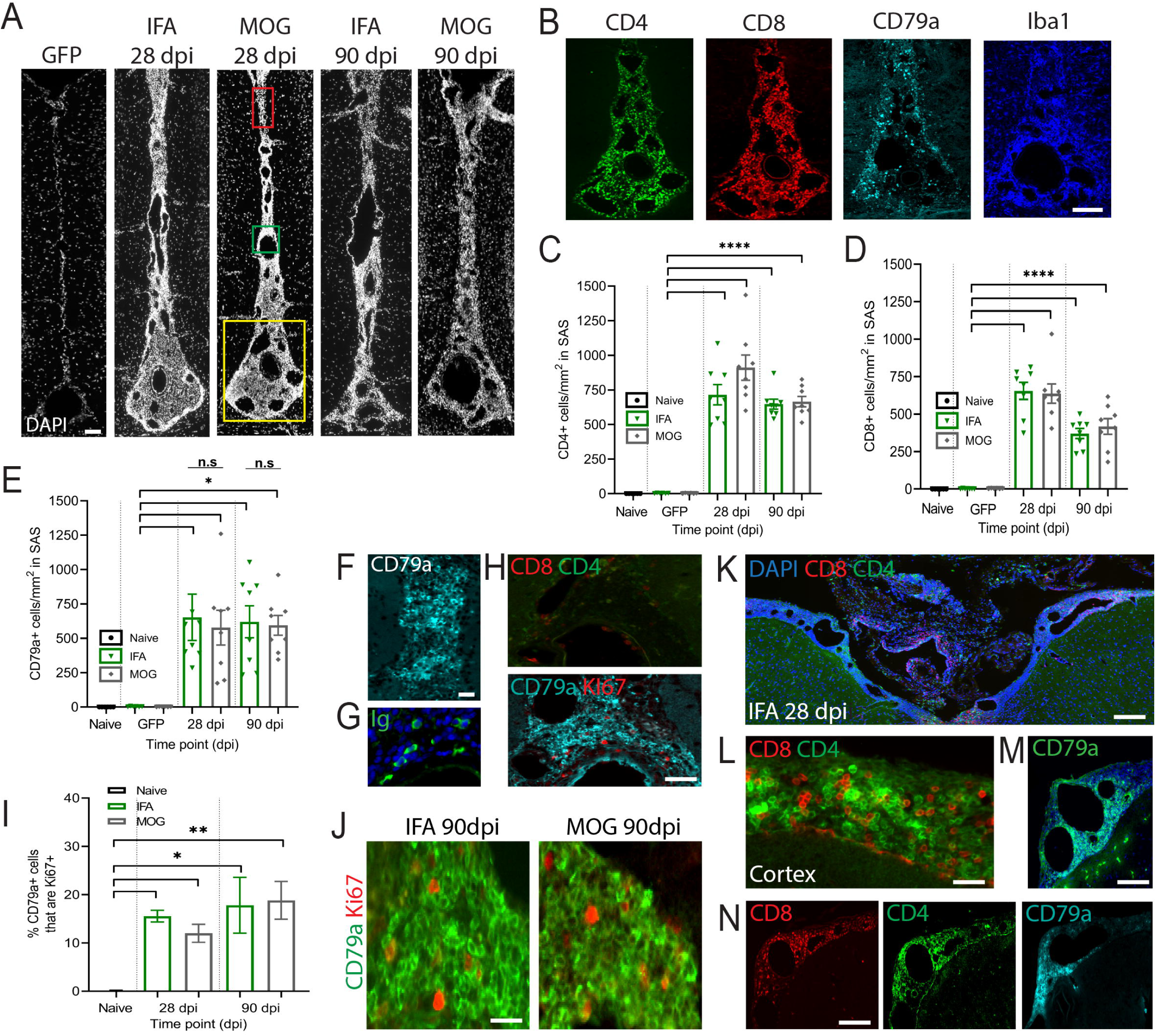
Overexpression of LTα in the subarachnoid space induces meningeal inflammation. **(A)** DAPI nuclear staining showing the accumulation of cells down the entire length of the sagittal sulcus in both IFA and MOG immunised animals at 28 and 90dpi. No accumulation of cells can be seen in GFP vector animals. Scale bar = 200µm **(B)** Immunostaining shows presence of CD4+ and CD8+ T-cells, CD79a+ B-cells and IBA1+ myeloid cells within the dense infiltrates. Scale bar =100µm. **(C-D)** Quantification of the numbers of CD4+ (C) and CD8+ (D) cells/mm2 down the length of the SAS in all animal groups. Data presented as mean ± SEM (n=3-4 per group). **** P<0.001 cytokine compared to naïve. **(E)** Quantification of CD79a+ B-cells expressed as cells/mm^2^ in the sagittal sulcus shows no difference in numbers between IFA and MOG immunised animals and between 28 and 90 dpi (* p<0.05). **(F)** Immunostaining of a cluster of CD79a cells within the SS. **(G)** Ig expressing plasma cells were seen within CD79a+ B-cell clusters. **(H)** Regions where there were dense clusters of CD79a+ cells showed very little presence of CD8 and CD4 cells, shown in a serial section. **(I)** The proportion of CD79a+ cells that were co-expressing the proliferation marker Ki67 **(J)** was similar for MOG and IFA immunised animals at both 28 and 90 dpi. **(K)** Dense aggregates of cells were also present across the surface of the cortex (scale bar=150µm), containing CD8+ and CD4+ T-cells **(L)** and clusters of CD79a+ B-cells (M), particularly around large channels or vessels. **(N)** CD4+ and CD8+ T-cells showed discrete localisation patterns that were distinct from CD79a+ cells. Statistics: 1-way analysis of variance with Tukey post-test. Scale bar H, M, N =100μm. Scale bar G,J,L = 40µm.

### Tertiary lymphoid structures in the meninges

The segregation of immune cells in the meninges into T-cell rich regions and large B-cell aggregates with ongoing proliferation was suggestive of TLS formation. Therefore, we further analysed the features of the areas of dense infiltration. A common feature was the presence of many channels of variable diameter, ranging from those the size of large venules (Fig. 6A, green arrows) to smaller sized channels reminiscent of high endothelial venules (HEVs; Fig. 6A, red arrows). Staining for the mucosal addressin cell adhesion molecule (MAdCAM-1) revealed multiple MAdCAM-1+ vessels throughout the length of the sagittal sulcus (Fig. 6B). MAdCAM-1 was expressed by large channels, on cells with the morphology of slender squamous endothelium cells (Fig. 6C) and on some process bearing individual cells (Fig. 6D). MAdCam-1 expression by mature stromal organiser cells has been previously reported (Benezech et al., 2010) which this population of cells may represent. Many of the channels were also strongly laminin positive (Fig. 6E), whereas the meningeal blood vessels were only weakly laminin positive. We were unable to further characterise the channels with lymphatic markers as all currently available commercial antibodies for mouse and human failed to react with the rat proteins. A dense matrix of stromal cells called fibroblastic reticular cells (FRC) are commonly found in the T-cell zone of TLS and express the markers podoplanin (gp38) and laminin. Podoplanin was highly expressed throughout the aggregates (Fig. 6F) particularly in regions that contained high numbers of CD4 and CD8+ T-cells. No difference in expression between IFA and MOG+IFA immunised animals was seen, with staining for a dense podoplanin network seen in both (Fig. 6G). No staining was present in naïve or GFP vector injected animals (not shown).

**Figure 6.**
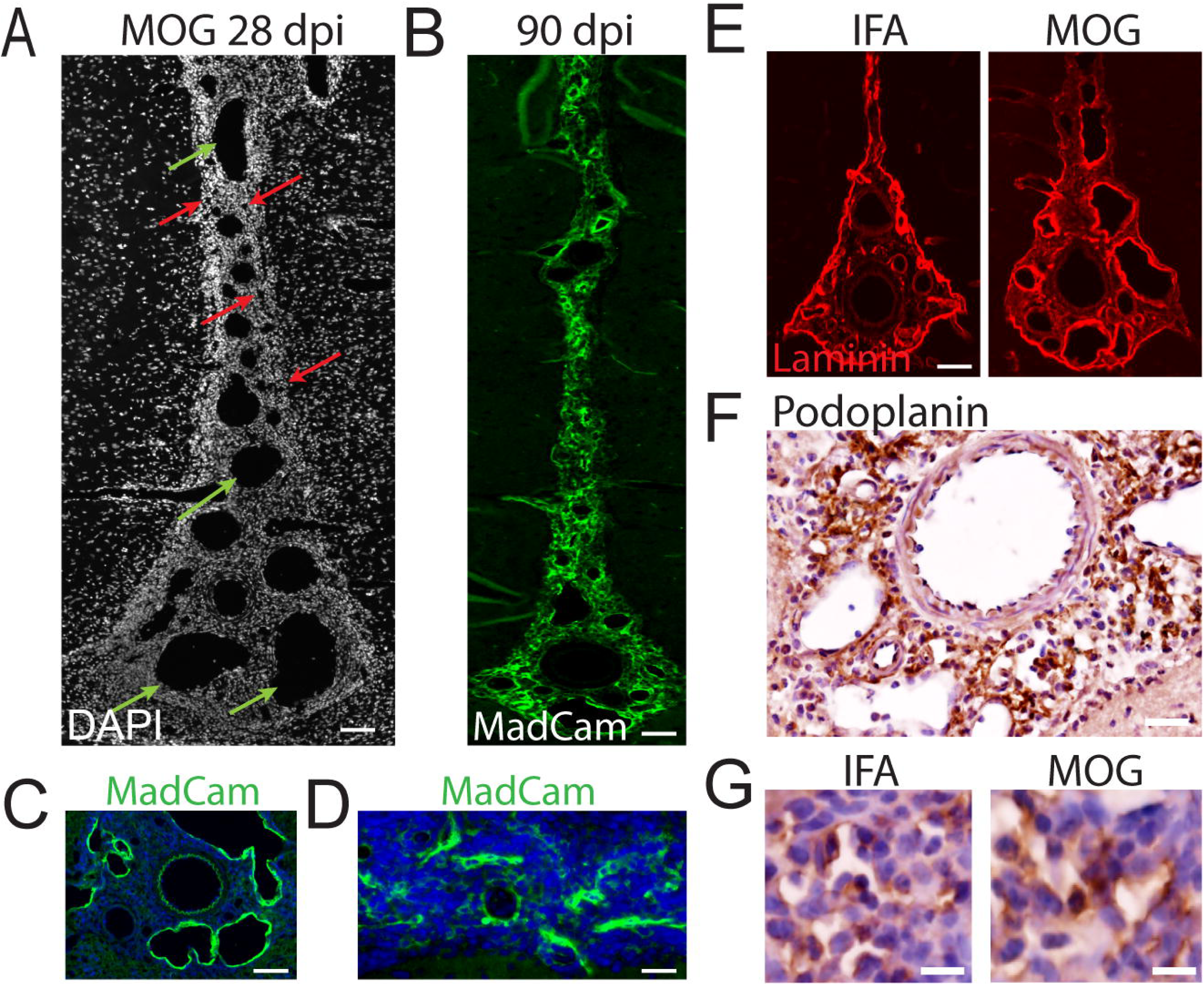
Presence of lymphatic channels in meningeal aggregates. **(A)** DAPI staining of meningeal aggregate at 28dpi in a MOG immunised animal showing the presence of multiple channels within the infiltrates that varied in size from small (red arrows) to very large, over 100µm in size (green arrows). **(B)** Immunostaining against MadCam-1 showed high expression the length of the SAS, particularly around the channels/vessels. Scale bar = 100um. **(C-D)** Higher magnification images show strong MadCaM immunofluorescence around large vessels/venules (C) as well as on a process bearing cellular network (D). **(E)** Laminin staining on the basement membranes surrounding SAS, illustrating the glia limitans boundary as well as around lymphatic vessels in both MOG and IFA immunised animals at 28 dpi. Laminin immunostaining on blood vessels was much weaker. **(F)** Podoplanin immunohistochemistry shows a reticulum of staining throughout the immune cell infiltrates. Scale bar =50µm. **(G)** Higher magnification images of podoplanin show IFA and MOG animals express similar staining patterns in the cytoplasm and processes of meningeal cells. Scale bar=20µm.

Binding of LTα homotrimers to the TNFR1/2 receptors and LTα/LTβ heterotrimers to the LTβR receptor induces expression of many chemokines involved in organisation of the lymphoid environment, many of which are increased in MS CSF (Magliozzi et al., 2018). To investigate which chemokines resident meningeal cells may produce locally, rat primary meningeal fibroblast cells were treated with 100ng/ml of recombinant rat LTα. LTα treatment induced significant increases in gene expression for CCL19, CXCL13 and LTβR after 24 hours (Fig. 7A). Transcript levels for BAFF, CXCL12 and CCL21 did not change significantly. RT-PCR on tissue dissected from the meninges of LVLTα animals at 28 dpi showed significant increases in the mRNA for CXCR5, CCL19, CCL21, CCR7, CXCR4, BAFF and MadCam compared to GFP injected animals (Fig. 7B). CXCL13 showed the largest increase in mRNA transcript levels (Fig. 7C). Immunostaining for CXCL13 revealed expression throughout the meningeal aggregates (Fig. 7D) that at higher magnification displayed an FDC-like fibroblastic expression pattern with multiple thin processes extending throughout the infiltrates (Fig. 7E). In regions where strong CXCL13 expression was found, a population of follicular dendritic cells (FDC) could be stained for the rat FDC marker ED5 (Fig. 7F). FDCs were typically found in B-cell rich areas, but the more widespread diffuse expression could explain why some CD79a+ cells were found distributed throughout the aggregates, even if in small numbers compared to T-cells (Fig. 5B). As expected, expression of the CXCL13 receptor, CXCR5, was found in close association within regions of high CXCL13 staining, along with clusters of CD79a+ B-cells (Fig. 7G).

**Figure 7.**
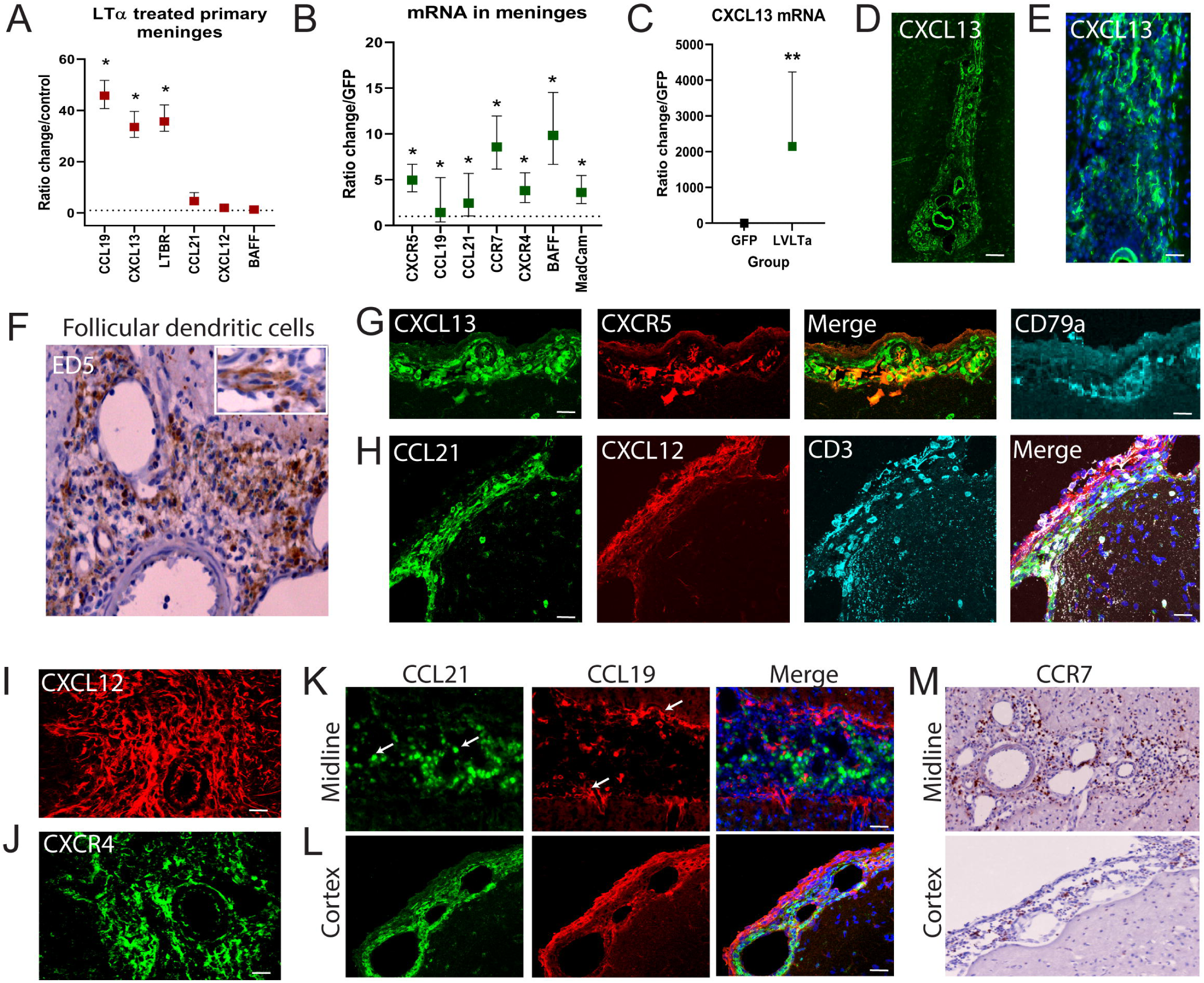
Expression lymphoid chemokines by meningeal cells. **(A)** Primary meningeal cells were treated with 100ng LTα for 24 hours and changes in gene expression measured by RT-PCR. Ratio changes calculated for LTα treated over non-treated showed increased gene expression levels for CCL19, CXCL13 and LTBR. * p<0.05. **(B)** The cortical meninges were microdissected from 4 x 10µm section from 28 dpi GFP or LVLTα injected animals for RNA extraction. Changes in mRNA levels for chemokines and receptors was measured by RT-PCR and shown here as Cytokine treated compared to GFP animals. * p<0.05 t-test on ΔΔCt values. **(C)** In the dissected meninges, CXCL13 transcript showed the greatest fold change in cytokine versus GFP animals at 28 dpi. ** p<0.01. **(D)** Immunostaining for CXCL13 at 28 dpi showed expression on blood vessels and lymphatic channels and on a network of cell processes through the immune cell infiltrate. Scale bar = 100µm. **(E)** Higher magnification shows CXCL13+ staining on cells with follicular dendritic cell morphology with multiple thin processes. Scale bar = 40µm. **(F)** IHC staining against ED5, a marker for follicular dendritic cells showed a dense network of cells through the infiltrates at 28 dpi. Inset box shows a higher magnification view. **(G)** CXCL13/CXCR5 double immunofluorescence and CD79a immunostaining on a serial section of cortical surface infiltrates. **(H)** Triple immunostaining for CCL21 and CXCL13 in regions with CD3+ T-cells showed distinct localisation patterns. **(I)** CXCL12 immunostaining was seen throughout the aggregates, together with expression of its receptor CXCR4 receptor **(J)**. CCL21 and CCL19 double immunofluorescence in midline **(K)** and cortical aggregates **(L)**. **(M)** Cells expressing CCR7 were found throughout the midline and cortical aggregates.

CXCL12+ expression, which provides a survival signal for incoming T-cells, was present on process bearing cells throughout the meningeal infiltrates (Fig. 7H and I) together with its receptor CXCR4 (Fig. 7J). Co-staining of CCL21 and CXCL12 in T-cell rich regions revealed that they were expressed by distinct populations of cells (Fig. 7H), with no apparent co-expression of the two chemokines. CCL19 and CCL21 also showed differential expression patterns with minimal overlap (Fig. 7K-L). CCL21 could be seen in rounded amoeboid-like cells surrounding large channels (Fig. 7K, white arrows green channel). In contrast, CCL19 was seen in more fibroblastic cells in the areas of infiltration along the edges of the sagittal sulcus (Fig. 7K, white arrow red channel) or cortical surface, in a pattern similar to that seen for CXCL12. Cells expressing the CCR7 receptor for CCL19 and CCL21 were present diffusely throughout the meningeal infiltrates (Fig. 7M).

### Demyelination and microglial activation after LTα overexpression

No observable demyelination was present at 28 or 90dpi following injection of LTα vector in animals immunised with only IFA (Fig. 8A-B). In contrast, extensive subpial demyelination was found in midline and cortical regions, which reached into deeper cortical layers, at both 28 and 90dpi in LTα vector injected animals immunised with MOG+IFA (Fig. 8A-B). Demyelination in MOG immunised animals was significantly greater at 90dpi than 28dpi in both midline (Fig. 8C; 41.0 ± 5.2% of cortical GM myelinated at 28 dpi versus 28.5 ± 4.6% at 90 dpi) and subcortical regions (Fig. 8D; 30.9 ± 2.0% myelinated at 28 dpi versus 17.8 ± 4.0% at 90dpi). GFP vector injected control animals showed no quantifiable demyelination, regardless of their immunisation status (Fig. 8C-D). Loss of MOG+ myelin extended from layer I directly beneath the pial surface down to layer III (Fig. 8E). In midline regions, loss of myelin extended from the midline into layer III (Fig. 8F). Loss of myelin basic protein (MBP) immunostaining confirmed the presence of demyelination rather than a loss of the MOG protein specifically (Fig. 8G). Microglial activation was extensive and widespread at 90dpi in both IFA only and IFA+MOG immunised LTα vector injected animals when compared to GFP vector injected controls (fig. 8H). In subpial cortical and midline regions, microglia showed a highly activated, ramified morphology, whilst in the deeper cortical layers V and VI directly above the corpus callosum they showed a more amoeboid morphology, (fig. 8I). In midline regions microglial numbers in LTα vector injected animals were increased at both 28 and 90 dpi when compared to naïve and GFP controls, with no significant difference between MOG+IFA versus IFA only immunised animals (Fig. 8J; 425 ± 37 cells/mm^2^ in IFA versus 507 ± 74 cells/mm^2^ in MOG animals at 90 dpi). In cortical layers IBA1+ cells were only significantly increased in MOG-immunised animals (Fig. 8K).

**Figure 8.**
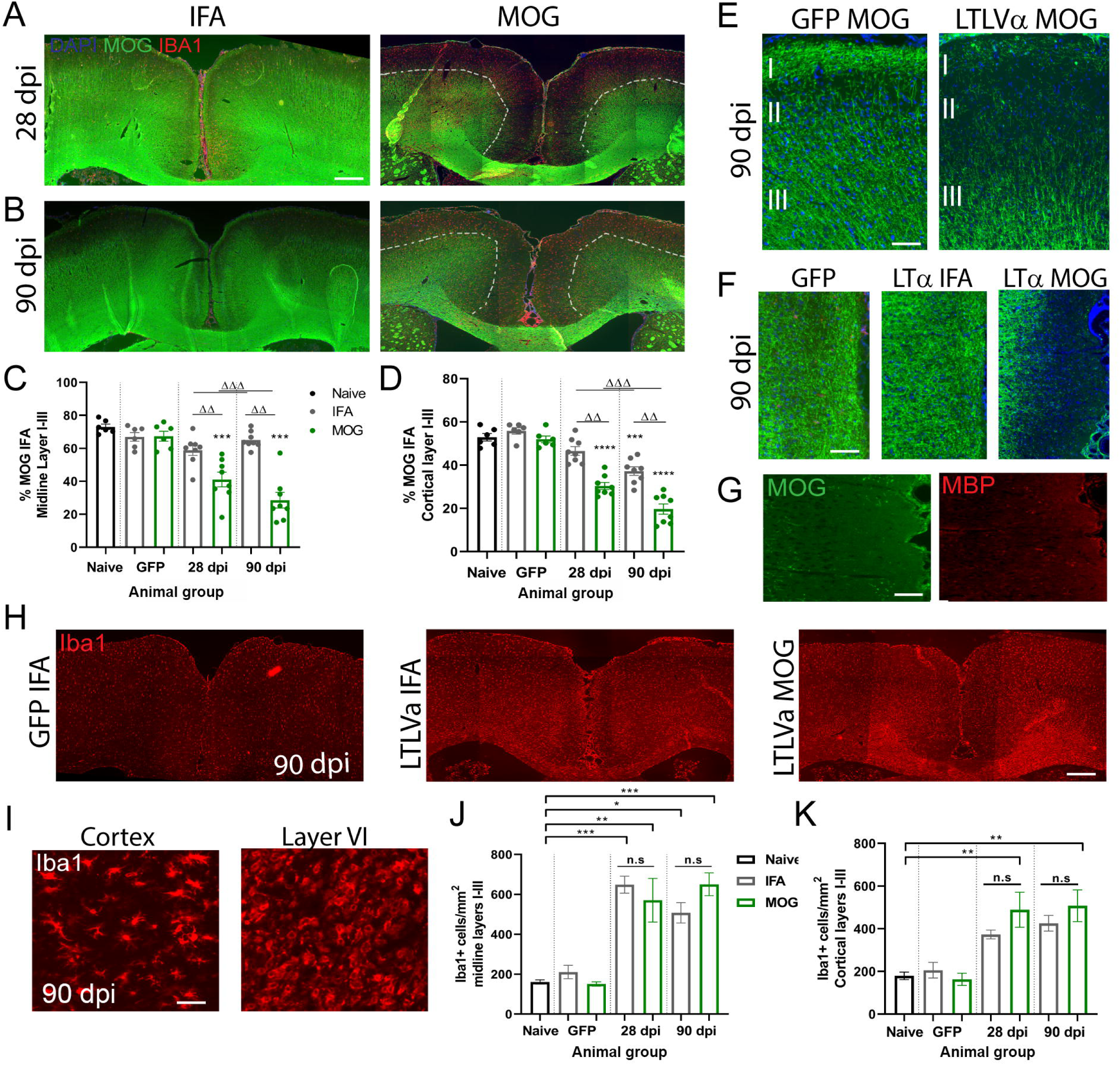
Demyelination after chronic overexpression of LTα. **(A-B)** Representative images of MOG staining in LVLTα injected animals immunised with IFA or MOG at 28dpi (A) and 90dpi (B). Demyelination was present in subpial areas in rmMOG immunised rats at 28 and 90 days after LVLTα injection (delineated by dotted lines). **(C-D)** Quantification of subpial demyelination in midline layers I-III (C) and cortical layers I-III (D) presented as the percentage area covered by MOG immunofluorescence (IF). Data is presented as the mean ± SEM. Statistics: 1-way analysis of variance with Tukey post-test. *** P<0.001; **** p<0.001 cytokine vs naïve. Naïve and GFP groups n=6, LVLTa groups n=8. **(E-F)** Immunofluorescence for MOG at higher magnification shows no demyelination in eGFP vector injected animals compared to MOG immunised animals in layers I and II in the cortex (E) or midline (F) at 90 dpi. **(G)** MBP immunostaining confirmed the lack of myelin shown by MOG immunostaining. **(H)** IBA1 immunofluorescence revealed little microglial activation immediately surrounding the sagittal sulcus and subarachnoid space in MOG immunised animals at 90 days post LV-GFP injection, whereas prominent microglial activation occurred throughout the corpus callosum and cortex following LVLTα injection. **(I)** Ameboid IBA1+ cells were found in layer VI compared to ramified IBA1+ cells in the surface cortex. **(J)** Quantification of IBA1 cells/mm^2^ showed significant increases in IBA1 cells/mm^2^ at 28 and 90 dpi in midline layers. **(K)** In the surface cortical regions only MOG immunised animals showed a significant increase in IBA1+ cell numbers. There was no significant difference between IFA or MOG immunised animals. * p<0.05, ** p<0.01, *** p<0.001. Statistics 1-way ANOVA with Tukey post test.

### Neuronal loss in cortical layers after persistent LTα overexpression

In order to determine whether chronic overexpression of LTα for 90 days would lead to cortical neurodegeneration, we counted neuronal numbers using NeuN immunostaining. In midline regions in close proximity to the meningeal aggregates there was a diffuse pattern of loss of neurons in layers I to III that extended the length of the sulcus (Fig. 9A). No neuronal loss was seen in GFP vector injected animals, suggesting this loss was not the result of the surgical procedure. Quantification of midline NeuN+ cell numbers showed a similar loss in both IFA (34.0 ± 2.3%) and MOG (34.0 ± 3.3%) immunised animals compared to both naive and GFP vector injected animals (Fig. 9B). In the cortical parenchyma away from the midline there were expansive regions throughout the motor and sensory cortices that demonstrated a marked decrease in NeuN+ cells, extending from the pial surface into the deeper cortical layers (Fig. 9C). Immunisation with MOG did not impact on neuronal loss (Fig. 9D). In cortical layers II-III there was a 40.0 ± 2.0% reduction in neuronal numbers in IFA and 33.0 ± 3.6% reduction in MOG immunised animals compared to naive. The cortical region with the most apparent loss of NeuN+ cells was the rostral cortex, where there were focal areas of loss, that in some animals extended from the subpial surface down to layer IV (Fig. 9E). This loss was apparent in all animals injected with LTα vector, both IFA and MOG immunised. We also found a significant reduction in the density of layer V neurons in the midline (Fig. 9F). Whilst there was some loss of larger pyramidal neurons, most of the loss seemed to represent smaller interneurons, although this distinction was not quantified (Fig. 9G). Examination of 200kDa neurofilament protein immunostaining in midline regions showed a decrease in staining in midline regions accompanying the decrease in neuronal numbers (Fig. 9H).

**Figure 9.**
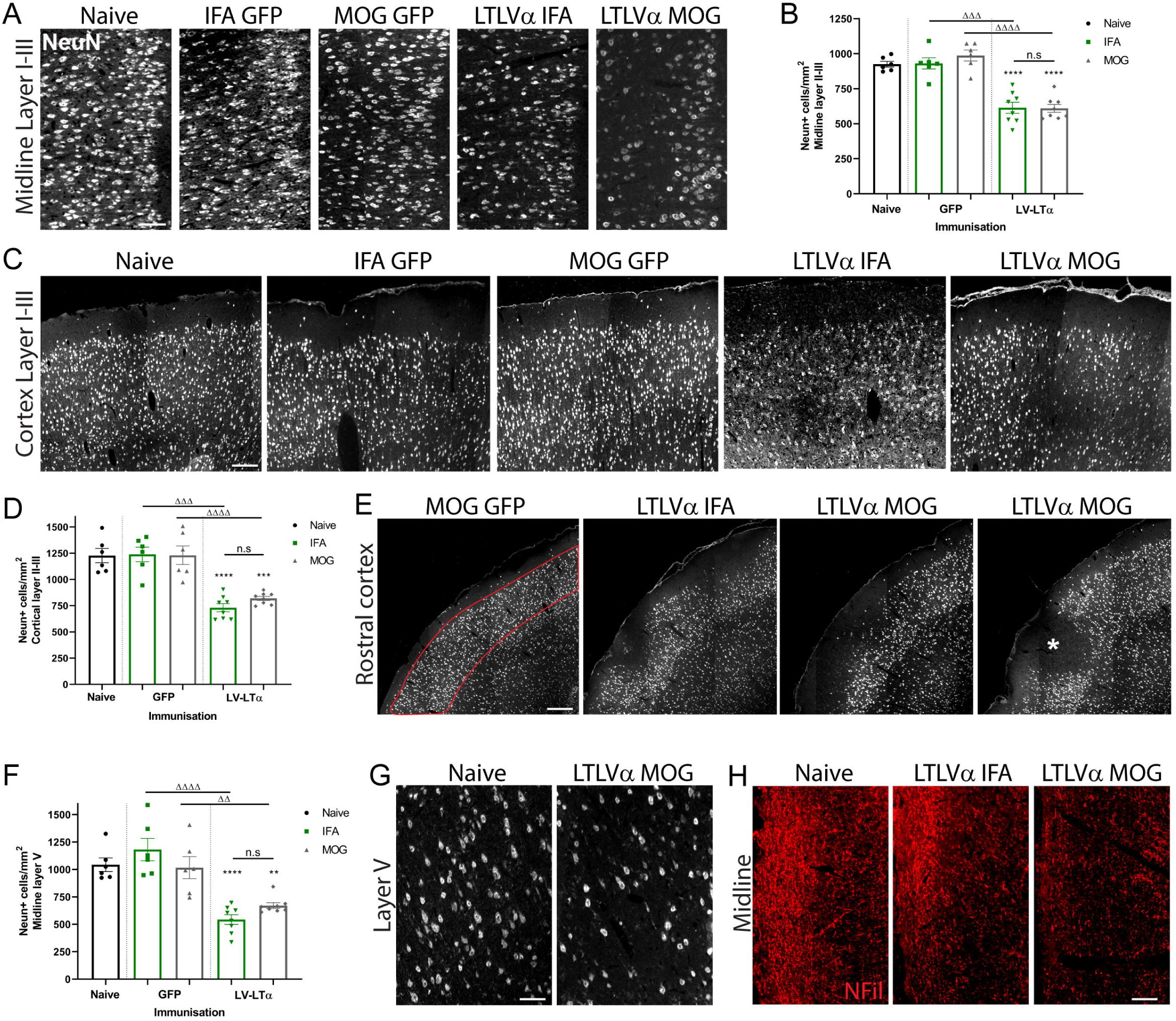
Neuronal loss in the cortical parenchyma after 90 days persistent LTα expression. **(A)** Representative images of NeuN immunofluorescence in midline layers I-III for naïve, LVGFP in IFA and MOG animals and LVLTα in IFA and MOG animals, showing a decreased neuronal density in LVLTα injected animals. Scale bar = 50μm. **(B)** Quantification of NeuN+ cells per mm^2^ in midline layer I-III regions after LV injection. GFP animals had no decrease in NeuN+ cells whilst LV-LTα animals had significant decreases in neuronal numbers both in IFA (613.8 cells/mm^2^, n=8) and MOG (609.6 cells/mm^2^, n=8) immunised animals compared to naïve (924.83cells/mm^2^, n=6). Graph presented is mean +/- SEM. ****p<0.0001. **(C)** Representative images of NeuN immunostaining in cortical layers I-III from motor cortex 90 days after cytokine or GFP vector injection. A decreased density of NeuN expressing neurons is seen in both IFA and MOG LV-LTα but not in GFP animals. Scale bar = 60µm. (D) Graph showing mean number of NeuN+ cells/mm^2^ in cortical layers I-III in all animal groups. A significant loss of NeuN+ cell is shown in both LVLTα injected IFA and MOG immunised animals when compared to GFP animals. There was no significant difference between IFA and MOG immunised LVLTα injected animals. **p<0.01, ***p<0.001, ****P<0.0001 cytokine versus naive. ΔΔ p<0.0, ΔΔΔΔ p<0.001 IFA /MOG versus IFA/MOG. (E) Representative images of NeuN immunostaining in rostral cortical regions. Loss of NeuN immunostaining in LVLTα animals compared to naïve controls was most visibly apparent in rostral regions, with loss present in both IFA and MOG animals. One MOG animal had focal neuronal loss that extended from the pial surface down through to layer IV (star in final image). Scale bar =100µm. (F) Quantification of NeuN+ cells/mm^2^ in cortical layer V. **p<0.01, ****p<0.0001 cytokine vs naïve. (G) NeuN+ staining in layer V of naïve and LVLTα MOG animals. In layer V there was a decrease in NeuN staining density, in particular for smaller size interneurons, rather than the larger pyramidal cells. (H) 200 kDa neurofilament protein immunofluorescence within the cortex in proximity to the midline showed a decreased density of staining in both IFA and MOG LVLTα animals compared to naive controls. Scale bar =50μm. Statistics - 1-way analysis of variance with Tukey post-test.

## Discussion

More extensive cortical demyelination and neurodegeneration in the MS brain, leading to an accelerated disease course (earlier age of clinical onset, more rapid disease progression and earlier age of death), has been shown to be associated with the presence of meningeal TLS formation (Howell et al., 2011; Magliozzi et al., 2007; Magliozzi et al., 2010; Serafini et al., 2004) and an increasingly pro-inflammatory CSF milieu (Magliozzi et al., 2018; 2020). In this study, we provide experimental evidence that chronic production of LTα in the cortical subarachnoid space of the adult rat can induce lymphoid chemokine expression, the persistent presence of meningeal immune cell infiltrates with many features of tertiary lymphoid structures, and chronic pathology in the underlying cortical GM that includes subpial demyelination, microglial activation and accumulating neuronal loss. Such findings are remarkably similar to the pathological changes seen in the MS cerebral cortex in SPMS and provide some possible insights into the pathogenetic mechanisms involved.

Numerous previous studies have shown the presence of LTα and β in serum and CSF from progressive MS patients and their production by activated T and B-cells cells (Buckle et al., 2003; Christensen et al., 2013; Corcione et al., 2004; Duddy et al., 2007; Matusevicius et al., 1996). In addition, the gene for LTα (*LTA* or *TNFSF1*) has been linked to MS susceptibility (Fernandes Filho et al., 2002) and is suggested to play a role in the stimulation of relapses in RRMS (Navikas et al., 1996; Rieckmann et al., 1995b). However, its role in the link between compartmentalised inflammation and the pathogenesis of CNS damage has received little attention. In our study, LTα post-mortem CSF levels were highest in patients that displayed the greatest levels of meningeal inflammation and the presence of meningeal TLS. Increased levels of LTα gene expression in the isolated meninges strongly suggests a cellular source, most likely from activated memory B-cells or CD4+ and CD8+ T-cells (Duddy et al., 2007; Yang et al., 2018). B-cells from untreated MS patients exhibit abnormal proinflammatory cytokine responses, in particular exaggerated production of LTα, when activated in the context of Th1 cytokine IFN1 or presence of TLR9 ligand (Bar-Or et al., 2010). Similarly, stimulation with the TLR9 agonist resulted in increased frequency of LTα producing B-cells in patients with RRMS (McWilliam et al., 2018). A significant increase in lymphotoxin secretion from anti-CD3-stimulated CD8+ T cells has also been reported in patients with SPMS compared to normal controls (Buckle et al., 2003), which is in agreement with the immunohistochemical localisation of LTα in CD3+ T-cells in the meningeal tissues in the current study. Therefore, it is possible that in MS cases with prominent meningeal inflammation, LTα expressed by infiltrating cells within the cerebral sulci may induce a sufficient increase in the local LTα concentration to result in fibroblast differentiation into FDCs and induction of lymphoid chemokines, including CXCL13, CCL19 and CCL21, as is suggested to occur in rheumatoid arthritis synovium (GeurtsvanKessel et al., 2009; Kain and Owens, 2013; Lindhout et al., 1999; van Nierop and de Groot, 2002).

Both acute and chronic elevation of LTα expression in the CSF were able to induce substantial immune cell infiltrates in the meninges that displayed similar cellular composition to the meningeal inflammation reported in SPMS patients (Bell et al., 2019; Magliozzi et al., 2010; Serafini et al., 2004), including those with short duration aggressive disease (Bevan et al., 2018). Although previous animal studies in both active and passive induction EAE models have described the presence of TLS at various stages of complexity in the meninges overlying the brainstem, cerebellum, spinal cord and in the ventricles (Kuerten et al., 2012; Magliozzi et al., 2004; Peters et al., 2011; Pikor et al., 2015), none of these models exhibited the cortical grey matter pathology that is commonly seen in MS. In addition, an association with neurodegeneration in the underlying neural tissue had not been examined. By creating a focal cortical model, with inflammation extending down into the full depth of the sagittal sulcus as well as across the cortical surface, we have reproduced a number of features of the situation seen in MS, notably the presence of TLS neogenesis in deep sulci (Griffiths et al., 2020; Howell et al., 2011). It is likely that the reduced CSF flow in the deep sulcus would allow a local increase in cytokine concentration that might not be reflected by global concentrations, although no experimental data is available on such CSF flow rates in the rat. In this regard it is also of interest that the EAE models in which TLS have been observed were characterised by a more chronic disease course (Kuerten et al., 2012; Magliozzi et al., 2004; Peters et al., 2011; Pikor et al., 2015), which would allow inflammatory infiltrates to build up and be maintained. Whilst TLS developed over a remarkably short time frame in the presence of acutely elevated LTα levels, it required persistent expression for them to be maintained. Persistent ectopic expression of LTα in peripheral organs in transgenic mice has been shown to lead to ectopic lymphoid neogenesis (Drayton et al., 2003; Kratz et al., 1996), but similar experiments have not investigated its role in the CNS and related tissues. The described model allows the investigation of some of the consequences of targeted increased LTα expression and subsequent compartmentalised inflammation on tissue damage in a way that is closely relevant to MS itself.

Persistent LTα expression might be expected to create a permissive microenvironment for TLS formation by endowing meningeal stromal support cells ‘immune competence’ to secrete lymphocyte chemoattractants, such as CXCL13, CCL19 and CCL21, thus producing support networks for the interaction of B-cells with T-cells and local activation and further maturation. Our data suggests that LTα may act on a population of meningeal follicular dendritic cells (FDCs) that then secrete CXCL13 and CCL19 in response to LTα stimulation, allowing the attraction and organisation of B-cells. A similar network of follicular dendritic-cells expressing CXCL13 has also been reported in TLS in the MS brain (Magliozzi et al, 2007; Serafini et al., 2004). It has been suggested that these FDCs are of stromal origin and could mature from ubiquitous local precursors such as PDGFRB+ pericytes in response to cytokines such as LTα (Krautler et al., 2012). Discrete patterns of staining also suggest the presence in the meninges of a population of stromal fibroblastic reticular-like cells (FRCs) that express podoplanin, CXCL12 and CCL21 and may help to form T-cell zones. A similar population of podoplanin+ FRCs that secrete CCL21, as well as CXCL13 and BAFF, has been previously reported in the mouse SJL/J EAE model (Pikor et al., 2015).

The presence of many channels of variable diameter within the dense meningeal infiltrates at 28 and 90 days post-LTα vector injection included some that resembled the “loose dilated” lymphatic vessels described previously in TLS in lung biopsies of idiopathic pulmonary arterial hypertension patients (Perros et al., 2012). Smaller channels with thin walls expressing laminin, podoplanin and MadCam-1, with an absence of red blood cells, suggests the formation of high endothelial vessels (HEVs) (Herzog et al., 2013). Unfortunately, expression of LYVE-1 and PNAd could not be used to confirm the presence of true lymphatic vessels as all the human and mouse reactive antibodies failed to react with the rat proteins. While few studies examining TLS in EAE have reported the presence of HEVs in the meninges (Cohen et al., 2021; Dang et al., 2015; Peters et al., 2011), HEVs expressing MadCam-1 and PNad have been reported in MP4-immunised mice (Kuerten et al., 2012). Given that HEVs develop in response to LTα it is possible their formation may only occur in the most mature TLOs or in cases with high LT levels (Browning et al., 2005). The meningeal aggregates seen in the current model bear much resemblance to TLS that develop in the kidney and pancreas of rat insulin promoter-LT (RIP-LTα/β) mice with regard to cellular composition (T cells, B cells, plasma cells, and antigen-presenting cells), delineated T and B cell areas and characteristic morphologic and antigenic (MAdCAM-1) features of high endothelial venules (Drayton et al., 2003; Kratz et al., 1996). This may indicate a very specific type of TLS that forms in the presence of LTα that may differ from those induced by other cytokines such as IL-17 (Pikor et al., 2015). However, further studies are required to elucidate the detailed cellular and molecular mechanisms by which they form in the cerebral meninges. Although our results show that LTα can play a major role in the initial formation of such structures, other cytokines and chemokines that have been demonstrated to be present in the MS meninges and CSF (Magliozzi et al, 2018) undoubtedly contribute.

Both acute and sustained LTα expression gave rise to subpial demyelinated lesions that extended from the pial surface down to cortical layer III/IV and demonstrated many similarities to key features of MS cortical lesion type III, in particular microglial activation, an absence of amoeboid macrophage-like cells and minimal lymphocytic infiltration. However, subpial demyelination required a pre-existing anti-MOG autoimmunity as very minimal demyelination occurred in IFA only immunised animals. The requirement for an anti-MOG response for demyelination to occur suggests that the microglial activation response may differ depending on prior immunisation, such that activated microglia release different effector molecules in rmMOG-immunised compared to IFA rats, or that autoantibodies contribute to microglial-mediated demyelination in this model. Autoantibodies may also induce demyelination through activation of complement, and immunoglobulin and complement fragment deposition have been observed on myelin sheaths at the lesion edge in targeted EAE following subarachnoid and intracortical injection of TNF and IFNγ (Gardner et al., 2013; Merkler et al., 2006) and in human MS cortical GMLs (Watkins et al., 2016), suggesting that humoral mechanisms may contribute to demyelination in these models and in the current study. In this scenario it is likely that LTα is mainly involved in targeting the immune response to the surface of the cortex, rather than playing a direct cytotoxic role in oligodendrocyte cell death, as has been reported *in vitro* (Selmaj et al., 1991b). The substantial areas of subpial demyelination that were observed in rmMOG-immunised rats at 28 and 90 days post-LVLTα, in contrast to the complete remyelination observed by 21 days after recombinant cytokine injection and in other targeted EAE models (Gardner et al., 2013; Merkler et al., 2006; Rodriguez et al., 2014), may represent chronic demyelination due to an eventual failure of remyelination, or overlapping cycles of demyelination and remyelination such that remyelination always appears incomplete. The pattern of demyelination suggests that complete failure ensues after longer time periods, although further studies are required to ascertain this.

Loss of neurons in the grey matter of the cerebral cortex of progressive MS brains, that is independent of demyelination, is a reproducible and widespread finding (Carassiti et al., 2018; Magliozzi et al., 2010; Picon et al., 2021; Trapp et al., 2018; Yates et al., 2017). It is more substantial in cases with higher levels of meningeal inflammation and lymphoid tissue formation, when it occurs in a decreasing gradient from the pial surface towards the white matter (Magliozzi et al., 2010). We report here a similar pattern of neuronal loss in midline and surface cortical regions following chronic expression of LTα in the overlying meninges, which is independent of demyelination or prior MOG immunisation. This result is similar to that obtained previously with TNF+IFNγ chronic expression (James et al., 2020), but in the case of LTα did not require the additional presence of IFNγ. Focal areas of neuronal loss were also found in the rostral cortex, including the somatosensory and secondary somatosensory cortex. Although the dense meningeal infiltrates did not spread this far across the cortex, it is possible that neurons in this region, which have been linked to walking and motor impairment in MS (Arpin et al., 2017a; Arpin et al., 2017b; Strik et al., 2020), are more susceptible to the proinflammatory environment created by overexpression of LTα in the CSF. Further work is needed to understand why these neurons might be particularly vulnerable.

The finding that neuronal loss was similar between MOG and IFA immunised animals and independent of demyelination suggests that LTα can be directly or indirectly cytotoxic to neurons. The LTα homotrimer, unlike the TNF homotrimer, only exists as a soluble protein and can bind to TNFR1, TNFR2 and HVEM receptors (Medvedev et al., 1996). Soluble TNF interaction with TNFR1 is known to predominantly stimulate cell death signaling, either apoptosis or necroptosis (Ofengeim and Yuan, 2013; Probert, 2015), whereas binding to TNFR2 is linked with tissue regeneration, including neuroprotection and remyelination (Madsen et al., 2016a; Madsen et al., 2016b; Raphael et al., 2019; Taoufik et al., 2011). Although constitutive expression of TNFR1 is low in the adult brain, it is significantly upregulated in MS and in the cortical GM is expressed predominantly on neurons (Magliozzi et al., 2019b; Picon et al., 2021). In the presence of increased meningeal inflammation and in a pro-inflammatory environment, TNF/TNFR1 mediated signalling in the cortical GM is directed towards necroptotic neuronal death (James et al., 2020; Magliozzi et al., 2019b; Picon et al., 2021). Therefore, it is possible that LTα could also directly stimulate neurodegeneration in the same way by binding to TNFR1, although more in vivo studies would be required to substantiate this. LTα can induce necroptosis with the same potency as TNF in vitro, and like TNF, utilises the RIPK1, RIPK3 and MLKL kinase cascade (Etemadi et al., 2013). The finding that CSF LTα levels were highest in MS patients with the greatest cortical thinning also supports the hypothesis that LTα could be acting directly on neurons following diffusion through the damaged pial membrane (Magliozzi et al., 2010). LTα could also be acting indirectly by stimulating chronic microglial and astroglial activation and subsequent release of neurotoxic mediators (Calabrese et al., 2015b). Increased levels of proinflammatory cytokines in the subarachnoid space have been shown to give rise to increased endogenous expression of TNF and IFNγ in the underlying cortical layers (James et al., 2020) and microglia in primary culture release glutamate in response to both LTα and TNF (Gallego-Delgado et al., 2020), raising the possibility that neurodegeneration could occur indirectly via excitotoxicity. However, further experimental studies are required to determine the detailed mechanisms by which LTα could be inducing neurodegeneration in this rat model. Whether stimulation of the formation of TLS in the meninges in this model is due to a direct effect mediated by LTα/TNFR1 interaction or by LTα monomers becoming incorporated into the cell membrane as LTa_2_b_1_ or LTa_1_b_2_ heterotrimers that subsequently bind to the LTβR, also remains to be determined.

In conclusion, we have demonstrated that overexpression of LTα in the meningeal space in rats leads to chronic cortical pathology that replicates many of the neuropathological and molecular characteristics of MS, including meningeal inflammation with lymphoid tissue formation, microglial activation, demyelination and neuronal loss. We show that chronic LTα overexpression alone is sufficient to induce and maintain meningeal TLS. The detailed molecular mechanisms that cause some MS patients to develop meningeal TLS and ways to reduce the formation of these meningeal immune cell infiltrates can now be tested through the use of therapeutic interventions using this experimental model.

## Materials and methods

### Human tissue sample selection and immunohistochemistry

Post-mortem brain tissue was provided by the UK Multiple Sclerosis Tissue Bank at Imperial College London, which obtains tissue through a prospective donor scheme, approved by the National Research Ethics Committee (80/MREC09/31). The diagnosis of all MS cases was confirmed by a consultant neuropathologist according to published guidelines, as described previously (Reynolds et al., 2011). Isolated meningeal tissue samples were dissected from 8 control cases with no neurological disease and 20 secondary progressive MS cases (Magliozzi et al., 2018). Details of each case are listed in Supplementary Table 1. Cases were classed as PM-MS high or PM-MSlow based on numbers of haematoxylin staining cell nuclei in 2 meningeal infiltrates and presence and extent of subpial grey matter demyelination, as described previously (Magliozzi et al., 2018). Protein levels of LTα were determined in post-mortem CSF obtained at autopsy from 20 SPMS, 10 with high (PM-MShigh) and 10 with low (PM-MSlow) meningeal inflammation, and 10 healthy controls, using immune-assay Luminex technology (Bio-Plex X200 System equipped with a magnetic workstation, BioRad, Hercules, CA, USA). All samples were run undiluted, in duplicate in the same experiment and in two consecutive experiments, in order to verify the reproducibility and consistency of the results as previously described (Magliozzi et al., 2018).

Naïve relapsing remitting MS patients were screened at diagnosis by 3T-MRI and stratified into MSHIgh (n=18) and MSLow (n=13) by the presence of at least 10 cortical lesions or less than 2 cortical lesions respectively (Supplementary table 2), as described for this patient cohort previously (Magliozzi et al., 2018). The control cohort (n=10) consisted of subjects with non-inflammatory neurological disease. CSF samples from patients were acquired at least 2 months from the last relapse, according to the Consensus guidelines for CSF and Blood biobanking. Samples were centrifugated before storage of the supernatant at -80⁰C until use. Levels of LTα were assessed using immune-assay multiplex techniques on the Luminex technology (Bio-Plex X200 system equipped with a magnetic workstation, BioRad, Hercules, CA), in the same way as the post-mortem samples.

### Quantitative reverse transcription polymerase chain reaction

Meningeal tissue was isolated by manual dissection using a scalpel from up to 16 snap-frozen cortical blocks per MS and control case to give a total mass of 100-250mg tissue per case (Gardner et al., 2013). RNA was extracted using an RNeasy Lipid Tissue Mini Kit (Qiagen) and quantified by Nanodrop ND1000 spectrophotometer. Reverse transcription of 1µg RNA per case to complimentary (cDNA) was performed using a Quantitect Reverse Transcription Kit (Qiagen) with integrated genomic DNA removal. Prime Time qPCR Assays (Integrated DNA Technologies, Iowa, USA) were used to quantify expression of LTα relative to the expression of two housekeeping genes (HKG; glyceraldehydes-3-phosphate dehydrogenase, GAPDH and Xpnpep1). Xpnpep1 is a stable HKG in post-mortem CNS tissue, with no known associations with neuroinflammatory or degenerative pathologies (Durrenberger et al., 2012). Reactions of 20µl were performed in triplicate for each case on a Stratagene MXP4000 system (Agilent Technologies), with MxPro 2007 software. An inter-plate calibrator did not detect any significant inter-plate differences in efficiency between successive runs. The thermal profile was 95° C for 10 min, 50 cycles with a two-step program (95° C for 15s, 60° C for 45s). Changes in genes expression were calculated using ΔΔCt method.

### Lentiviral vector production

Lentiviral (LV) vectors expressing human LTα (LVLTα) or enhanced green fluorescent protein (LVGFP) were produced as described previously (James et al., 2020). We utilised a HIV-1 transfer plasmid (326-pRRL-sincppt-CMV-eGFP-WPRE genome plasmid) that carries the human cytomegalovirus promoter (CMV) for lentiviral production. Complementary DNA sequences for human lymphotoxin alpha (LTα) were codon optimised for rat including a 5’ Kozak sequence and synthesised by Gene Art with Xba1 and Sal1 restriction sites (Life Sciences, Paisley, UK). The human transgene DNA fragments were excised by restriction digest with Xba1 and Sal1 before purification using preparative agarose gel electrophoresis. The eGFP was removed from the transfer plasmid by digestion with Xba1 and Sal1 and the transgene DNA fragments ligated in-frame using the same restriction sites.

Recombinant HIV-1 based lentiviruses were produced using four plasmid co-transfection of HEK-293T cells as described previously (James et al., 2020). Briefly, HEK-293T cells were transfected with 15μg vector plasmid (pRRLsincppt-CMV-LTα-WPRE or pRRLsincppt-CMV-GFP-WPRE), 15μg of the packaging vector plasmids expressing the HIV-1 gag/pol gene (pMD2-LgRRE), 3μg of HIV-1 Rev (pRSV-Rev) and 5.1μg VSV-G envelope plasmids (pMD2-VSV-G) following the addition of 2 mol/L CaCl_2_. Lentivirus was concentrated from supernatant using ultracentrifugation and the genome copy number was calculated using the Clontech Lenti-X qRT-PCR Titration kit (Takara).

### Transduction of HEK cells

Hek-293T cells, obtained from ATCC, were maintained at 37C, 5% CO2 environment and cultured in Dulbecco’s modified Eagle’s medium (Sigma, UK), supplemented with 10% Newborn Calf serum (Heat inactivated, Sigma UK), 2 mM penicillin/streptomycin and 2 mM L-glutamine (Sigma, UK). Hek-293T cells were plated in 24 well plates for 24 hours before transduction with LV-LTα or LV-GFP at MOI 5, 10, 20, 35 and 50 and the supernatant harvested after 24 and 48 hours.

### Transduction of primary meningeal cultures

Primary meningeal cultures were prepared from P4 Sprague Dawley pups. Briefly, the meninges were dissected from the surface of the cortex and digested with papain and DNAse. Cells were plated on poly-l-lysine coated 8-well chamber slides or 24 well plates (Corning) and grown in DMEM/F12 (Sigma, UK) supplemented with 10% calf serum, 1% Penicillin/Streptomycin and 1% L-Glutamine (Sigma, UK). After 1 week in culture cells were transduced at various MOIs with LV-LTα, or LV-GFP and the supernatant harvested after 72 and 96 hours and 7 days. Enzyme-linked immunosorbent assay for human LTα was performed in triplicate using the DUO set ELISA kit (DY210, R&D Systems, Abingdon, UK). Protein was extracted from transfected meningeal cells using RIPA buffer (R0278, Sigma) and run on Western blot at 20ug per well. Protein was stained with an antibody specific for human LTα to confirm expression.

For gene expression analysis of meningeal cells, 1×10^5^ cells were plated in 6 wells plates and treated with 100ng recombinant human LTα. RNA was extracted using TRIzol reagent according to manufacturers instructions and 1μl of RNA converted to cDNA using High-capacity cDNA reverse transcription kit (Thermofisher, Uk). Prime Time qPCR Assays (Integrated DNA Technologies, Iowa, USA) were used to quantify expression of LTα relative to the expression of two housekeeping genes (CYC1 and TOP1). Reactions of 20µl in PowerUp SYBR green (Thermofisher, UK) were performed in triplicate for each case on a Stratagene MXP4000 system (Agilent Technologies), with MxPro 2007 software. The thermal profile was 95° C for 10 min, 40 cycles with a two-step program (95° C for 5s, 60° C for 30s). Changes in genes expression were calculated using ΔΔCt method.

### Immunofluorescence of primary meningeal cells

For immunofluorescence of transfected meningeal cells grown in 8-well chamber slides, cells were fixed with cold 4% PFA for 1 hour at room temperature. After fixation, cells were incubated with 0.1M PBS-0.1% triton-X-100 for 1 hour. Primary antibodies were added at their corresponding working concentration for 1 hour at room temperature. After primary antibody incubation, cells were thoroughly rinsed in 0.1M PBS, three times 10 min each. After rinsing cells with incubated with appropriate secondary antibody conjugated to the appropriate fluorophore for 1 hours at room temperature. The cells were rinsed in 0.1M PBS (three times, 10 min each) and dH_2_O, nuclear counterstained with DAPI (diluted in 1:10,000). Lastly, the tissues were mounted with Vectashield Antifade Mountant and coverslipped (24×50mm, VWR, International).

### Animals

All *in vivo* procedures were performed under UK Home Office approval. Eight to ten week old female Dark Agouti (DA) rats (140-160g) were obtained from Janvier (France) and kept in groups of four in a 12h light/dark cycle with food and water provided ad libitum.

### Induction of sub-clinical experimental autoimmune encephalomyelitis

Sub-clinical MOG-induced EAE was induced as described previously (Gardner et al., 2013; James et al, 2020). Rats were anaesthetised with isofluorane and immunised intradermally at the base of the tail (dorsal aspect) with 5μg of recombinant mouse MOG (amino acids 1-119, corresponding to the external Ig-like domain) diluted in sterile PBS and emulsified in an equal volume of incomplete Freund’s adjuvant (IFA, Sigma; total volume 100µl per rat). IFA control rats were immunised with PBS emulsified in an equal volume of IFA, and naïve age-matched control rats (n=6) received no treatment. For all groups n was 5 to 7 animals with a total of 92 animals used for all experiments. All animals were weighed, examined and clinically scored daily according to a scoring system to define the level of neurological deficit, as described previously (Papadopoulos et al., 2006). This dose of recombinant MOG protein has been previously shown to elicit only sub-clinical EAE in DA rats of similar age and weight (Gardner et al., 2013), with production of peripheral anti MOG IgG1 and IgG2a antibody titres previously confirmed to be present in MOG immunised but not IFA immunised animals (James et al., 2020).

### Subarachnoid injection of cytokines and LV vectors

Following immunisation, stereotaxic injections of recombinant cytokines or lentiviral (LV) vectors into the subarachnoid space (SS) were performed under isofluorane anaesthesia at 20-23 days post immunisation, or at an equivalent time-point in age-matched naïve control animals. A 5mm hole was drilled in the skull at 0.9mm caudal to bregma at the level of motor cortex. A finely calibrated pulled glass capillary needle attached to a 26S fixed needle Hamilton syringe (10μl) was lowered to a depth of 2.3mm ventral to the dural membrane. The cytokines or lentiviral vectors were infused at a rate of 0.2µl/min over a period of 20 minutes. Monastral blue (copper II phthalocyanine-tetrasulfonic acid tetrasodium salt; Sigma-Aldrich) was added as a tracer to determine the location of the injections, and resulted in no pathology, as confirmed by previous studies (Kerschensteiner et al., 2004; Merkler et al., 2006; Rodriguez et al., 2014). For lentiviral injections, 4μl of lentiviral vector solution diluted in TSSM with 0.5mM monastral blue at a total of 5 x 10^8^ genomic copies (GC) for LTα and GFP was injected at a rate of 0.2μl/min. For recombinant cytokine injections, control animals received an injection of sterile PBS 0.5% BSA (n=5). Recombinant human lymphotoxin-alpha (hLTα, reconstituted in 0.1% (w/v) BSA in sterile PBS; Peprotech) and recombinant rat IFNγ (reconstituted in pH 8 sodium phosphate [10mM; Sigma-Aldrich] were diluted to the required concentrations (LTα 1μg + IFNγ 75ng per injection) and injected at a total volume 2μl per rat. For all injections, the glass capillary needle was left in place for 10 min after the injection to allow diffusion of the injected solution away from the injection site before slow retraction of the capillary. A group of age-matched naïve rats, which received no immunisation or SS injections, provided an additional control (n=6).

### Tissue processing and CSF analysis

At the termination of the experiment rats were killed by intraperitoneal injection of 200 mg/kg of sodium pentobarbitone and a 100µl CSF sample taken from the cisterna magna using a 26G needle (Hamilton) before cessation of breathing, with the animal on a stereotaxic frame. Animals were perfused with 100ml of PBS followed by 400ml of 4% PFA through the left ventricle at 3, 7, 14, 21, 28 and 56 days post viral injection. Brains were removed and post fixed in 4 % PFA (4 hrs room temp), prior to cryoprotection in 30% sucrose solution (w/v in PBS) for 72 hours until equilibrated. Brains were embedded in optimal cutting temperature compound (OCT; Tissue-Tek, Sakura, Netherlands) and frozen in isopentane on dry ice. Coronal cryosections (10μm) were stored at -20C prior to performing immunofluorescence. Protein expression for the expressed human LTα transgene was measured in snap-frozen rat CSF using a human TNF-beta/TNFSF1 Quantikine ELISA kit (DTB00, R&D systems) as per manufacturer’s instructions.

### Immunohistochemistry and immunofluorescence

Immunofluorescence was performed as previously described (James et al., 2020). For immunohistochemistry snap-frozen sections sections were fixed with cold-methanol (4° C) for 10 minutes and blocked with 10% normal horse serum (NHS), followed by overnight incubation with primary antibody, and then incubated with ImmPRESS HRP-conjugated secondary antibodies (Vector laboratories). For single IHC, slides were visualised with ImmPACT-DAB (Vector laboratories) as the chromagen. Dual colour IHC was performed by sequential staining, followed by detection with ImmPACT-DAB, incubation with the second primary antibody and detected using the ABC-alkaline phosphatase detection system with Vector blue as substrate. The primary antibodies used were: mouse anti-MOG (clone Y10, Prof Reynolds, Imperial College London, UK); rabbit anti-myelin basic protein (MBP) (Polyclonal, Merck, Darmstadt, Germany); mouse anti-neurofilament-H protein (clone NE14; Merck, Darmstadt, Germany); rabbit anti-IBA1 (Polyclonal IgG, Wako Pure Chemical Corporation, USA); rabbit anti-Glial Fibrillary Acidic Protein (GFAP) (Polyclonal, Dako Agilent, Santa Clara, CA, USA); mouse anti-NeuN (clone A60, Merck, Darmstadt, Germany); mouse anti-CD79a (ThermoFisher); mouse anti-CD4 (AbD Serotec); mouse anti-CD8 (AbD Serotec) goat anti-human LTA (R&D systems); rabbit anti-laminin (Wako, Japan); rabbit anti-cleaved caspase-3 (Cell signalling, USA); goat anti-CXCL13 (R&D systems); mouse anti-CCL21 (R&D systems); mouse anti-CCL19 (R&D systems); mouse anti-CXCL12 (monoclonal, Sigma UK); rabbit anti-CXCR4 (Sigma, UK); goat anti-LTBR (polyclonal Goat, R&D); rabbit anti-Ki67 (Santa-Cruz); rabbit anti-CD3 (DAKO); rabbit anti-IgG (R&D); rabbit anti-laminin (Novusbio); mouse anti-MadCam (monoclonal mouse, R&D systems); mouser anti-CNPase (Millipore); rabbit anti-CXCR5 (Abcam, UK), rabbit anti-podoplanin (Abcam, UK); mouse anti-CCR7 (mouse monoclonal, R&D systems).

All the secondary antibodies used for immunofluorescence were purchased from ThermoFisher Scientific (USA): Alexa Fluor 546 Goat Anti-Mouse IgG (H+L), Alexa Fluor 546 Goat Anti-Rabbit IgG(H+L), Alexa Fluor 488 Goat Anti-Mouse IgG (H+L) Fluor 488 Goat Anti-Rabbit IgG (H+L), Alexa Fluor 488 Goat Anti-Mouse IgG1, Alexa Fluor 546 Goat Anti-Mouse IgG1, Alexa Fluor 647 Goat Anti-Mouse IgG1, Alexa Fluor 488 Goat Anti-Mouse IgG2b, Alexa Fluor 647 Goat Anti-Mouse IgG2b, Alexa Fluor 488 Streptavidin, Alexa Fluor 546 Streptavidin, Alexa Fluor 647 Streptavidin. To couple with Alexa Fluor labelled Streptavidin, the following biotinylated antibodies were used from Vector Laboratories (UK): biotinylated goat anti-rabbit IgG (H+L), biotinylated horse anti-mouse IgG (H+L) and biotinylated goat anti-mouse IgG2a (Life Technologies, ThermoFisher Scientific).

### Image acquisition

Immunostained sections were viewed with either an epifluorescence Olympus BX63 scanning microscope or a SP8 Leica confocal microscope for the rat sections and a Zeiss Axioscope 5 for the human sections. Tiled digital images were obtained at x10 or x20 magnification and coded for blinded analysis using ImageJ software (NIH, Maryland, USA). Higher magnification images were acquired on Leica Confocal at 60x magnification in 8μm z-stacks.

For each animal, immunofluorescence tiled images were acquired from 4 x 10μm sections stained with anti-MOG antibody to quantify demyelination. Cortical and midline layers were delineated as regions of interest (ROI) and automated Otsu thresholding was used to calculate the percentage area of ROI stained by the MOG antibody. For recombinant protein study thresholding was also used to calculate the area of IBA1 immunoreactivity (IR). For quantification of meningeal inflammation images of the length of the sagittal sulcus were acquired at 20x. DAPI staining was used to calculate the area of the subarachnoid space from the base of the sagittal sulcus to the position of the lateral blood vessels visible in the meninges. Meningeal immune cell numbers were counted manually for CD4, CD8 and CD79a expressing cells using the cell counting tool in ImageJ software and expressed as cells/mm^2^, from 4 x 10μm sequential sections spanning a 80μm distance in the anterior/posterior plane.

For both neuronal and microglial numbers, regions of interest were drawn that outlined the cortical layers or midline regions using ImageJ. Cortical layers were determined using co-staining with DAPI and GluR2/3 to identify different cortical regions (Gardner et al., 2013; James et al., 2020). Cortical regions began from the lateral vessels in the meninges at the top of the sagittal sulcus and extended the breadth of the cortex to the rostral regions. NeuN+ and IBA1+ cell numbers were counted manually at 20x magnification using the ImageJ cell counter tool. The total number of cells for all the counts was then divided by the total area of the region of interest to give the total cells/mm^2^. For neurons and microglia, the number of cells was calculated from four sections per animal that were spaced 10μm apart covering a total distance of 80μm per animal.

### Statistical analysis

Graphpad Prism8 statistical software (La Jolla, CA, USA) was used in all cases to present the data and to conduct statistical analysis. All data on graphs are expressed as mean ± SEM except for PCR data which is shown as ΔΔCT with standard errors. Group comparisons for cell quantifications were analysed by one-way ANOVA with Tukey test for multiple comparisons.

## Online supplemental material

Fig. S1 shows the testing and efficiency of the lentiviral vectors expressing human LTα eGFP following transduction by primary rat meningeal cells in culture. Table S1 lists the demographics and clinical characteristics of the post-mortem MS and control cases used for CSF and tissue analysis. Table S2 lists the demographics and clinical characteristics of the MS and control patient cohorts used for CSF analysis, as previously presented in Magliozzi et al., 2018.

## Acknowledgements

The authors would like to thank the Central Biomedical Services Staff at Imperial College for their help with animal welfare. This work was supported by the Multiple Sclerosis Society of Great Britain (grant 978/12 to RR and NDM and 037/15 to RR and REJ). NDM was supported by the European Research Council (7th Framework Proof of Concept grant no. 620253). EB and RS were supported by a PhD studentship from the UK Medical Research Council.

## Author contributions

R. James contributed to the concept and design, acquisition, analysis and interpretations of the data and drafting of manuscript. E. Browne contributed to concept and design, acquisition, analysis and interpretations of the data and drafting of manuscript. R. Schalks contributed to concept and design, acquisition and analysis of data. L. Tan, P. Parakesh and H. Jacobs contributed to acquisition and analysis of data. R. Magliozzi contributed to analysis of human tissues and revision of the manuscript. M. Calabrese contributed to the clinical analysis of the patient cohort. N.D. Mazarakis contributed to concept and design, obtaining research grants and revision of the manuscript. R. Reynolds contributed to concept and design, acquisition, obtaining research grants, interpretation of data and drafting and revision of the manuscript.

---

**Supplementary figure 1. Primary meningeal cells transduced with lentiviral vectors expressing human LTα.** Primary meningeal cells were produced from meninges dissected from P2 rat pup cortices and cultured for 3 weeks. **(A)** Around 20% of cells expressed TNFR1 and **(B)** approximately 98% of cells expressed laminin. **(C)** Cells transduced with lentiviral vectors (LVs) expressing enhanced GFP (LVGFP) did not any obvious changes in cell morphology or signs of stress and show high levels of GFP expression at 72hours following transduction **(D)**. **(E)** Immunostaining untreated control meningeal cells with an antibody specific to the human form of lymphotoxin alpha showed no constitutive expression. **(F)** Following transduction with LV-LTα at an MOI of 1000, immunostaining with a hLTα antibody showed a transduction efficiency of around 98% 72 hours post-transduction. Cleaved caspase-3 staining showed apoptotic cells were rare **(G)**. **(H)** LTα expression by human embryonic kidney 293T (HEK) cells at 24 **(H)** or 48 **(J)** hours after transduction with 5,10, 20, 35 or 50 MOI was determined by ELISA. LTα supernatant concentration was significantly increased following transduction with 35 and 50 MOI LVLTα compared to control cells (medium only) and LVGFP (p=0.0422 and 0.0059 respectively, Kruskal-Wallis with Dunn’s post-test**). (K)** Transduction of passaged meningeal cells with lentiviral vectors following longer term culture. LTα supernatant concentration was significantly increased at 7 days post-transduction of meningeal cells with 2000 MOI LVLTα compared to control or LVGFP. **(L)** A western blot image of meningeal cell lysates at 72 hours post-LVGFP or LVLTα transduction showed a 42kDa band corresponding to β-actin in all lanes and a 22kDa band corresponding to human LTα post LVLTA but not LVGFP transduction. Statistics - 1-way analysis of variance with Tukey post-test.

**Supplementary Table 1:** Demographic and clinical characteristics of the post-mortem cases used for CSF analysis and immunohistochemistry

| Cases | Sex | Age at Death | Cause of death | PM interval (hrs) | Disease duration (yrs) |
| --- | --- | --- | --- | --- | --- |
| **Control cases** |  |  |  |  |  |
| C5 | F | 95 | Bronchopneumonia | 9 | - |
| C7 | F | 85 | Esophageal cancer | 9 | - |
| C14 | M | 64 | Cardiac failure | 18 | - |
| C28 | F | 60 | Ovarian cancer | 13 | - |
| C30 | M | 75 | Bronchopneumonia | 17 | - |
| C32 | M | 88 | Prostrate cancer | 12 | - |
| C36 | M | 68 | Cardiac failure | 30 | - |
| C45 | M | 77 | Cardiac failure | 22 | - |
| C48 | M | 68 | Colon cancer | 10 | - |
| C50 | M | 32 | Haemangiopericytoma, | 6 | - |
| C51 | M | 68 | Ischemic heart disease | 24 | - |
| C54 | M | 66 | Pancreatic Cancer | 16 | - |
| PDC8 | F | 71 | Myocardial infarction | NA | - |
| PDC22 | M | 75 | Lung carcinoma | 10 | - |
| PDC32 | F | 91 | Colon cancer | 12 | - |
| PDC36 | F | 57 | Breast cancer | 15 | - |
| **MS cases - low meningeal inflammation** |  |  |  |  |  |
| MS104 | M | 53 | Multiple sclerosis | 12 | 11 |
| MS296 | M | 59 | Multiple sclerosis | 22 | 40 |
| MS301 | F | 62 | Septicaemia, MS | 16 | 20 |
| MS304 | M | 52 | Pulmonary embolism | 13 | 23 |
| MS311 | F | 45 | Pneumonia | 22 | 17 |
| MS318 | F | 59 | Multiple sclerosis | 13 | 34 |
| MS326 | M | 62 | Prostate cancer, MS | 24 | 32 |
| MS335 | M | 62 | Renal failure | 22 | 38 |
| MS347 | M | 50 | Pancreatic carcinoma | 13 | 28 |
| MS364 | F | 56 | Bronchopneumonia | 14 | 34 |
| MS376 | F | 58 | Multiple sclerosis | 19 | 21 |
| MS422 | M | 58 | Bronchopneumonia | 25 | 13 |
| MS444 | M | 49 | Renal failure | 18 | 20 |
| MS461 | M | 43 | Bronchopneumonia | 13 | 21 |
| MS485 | F | 57 | Bronchopneumonia | 24 | 29 |
| **MS cases - high meningeal inflammation** |  |  |  |  |  |
| MS180 | F | 44 | Multiple sclerosis | 9 | 18 |
| MS234 | F | 39 | Aspiration pneumonia | 15 | 15 |
| MS289 | M | 45 | Multiple sclerosis | 9 | 18 |
| MS317 | F | 48 | Aspiration pneumonia | 21 | 30 |
| MS330 | F | 59 | Bronchopneumonia | 21 | 40 |
| MS352 | M | 43 | Bronchopneumonia | 26 | 19 |
| MS356 | F | 45 | Multiple sclerosis | 10 | 16 |
| MS371 | M | 40 | Bronchopneumonia | 27 | 17 |
| MS377 | F | 50 | Aspiration pneumonia | 22 | 23 |
| MS402 | M | 46 | Multiple sclerosis | 12 | 20 |
| MS407 | F | 44 | Septicaemia | 22 | 19 |
| MS408 | M | 39 | Pneumonia | 21 | 10 |
| MS423 | F | 54 | Pneumonia | 11 | 30 |
| MS426 | F | 48 | Pneumonia | 21 | 19 |
| MS438 | F | 53 | Multiple sclerosis | 17 | 18 |
| MS473 | F | 40 | Broncho pneumonia | 9 | 13 |
| MS497 | F | 60 | Aspiration pneumonia | 26 | 29 |
| MS513 | M | 51 | Multiple sclerosis | 17 | 18 |
| MS517 | F | 48 | Pulmonary sepsis | 12 | 25 |
| MS528 | F | 45 | Multiple sclerosis | 17 | 25 |

Details of the control cases, and MS cases. M = male; F = female; PMD = post-mortem delay; yrs = years; NA = information not available. Cause of death is the primary cause taken from the certificate of death.

**Supplementary Table 2:** Demographics and disease characteristics of the MS subjects

| Group | Naïve MS patients |  |
| --- | --- | --- |
|  | Low (13) | High (18) |
| Age studied (mean±SD) | 38.4±11.3 | 33.2±11.1 |
| Gender (f:m) | 8:5 | 13:5 |
| Disease duration (mean±SD years) | 4.1±2.4 | 5.3±2.0 |
| EDSS at recruitment (range) | 1.5 (1.0-3.5) | 2.0 (1.0-4.0) |
| CLs volume (mm<sup>3</sup>)(range) | 49 ± 74<br>(0-188) | 1713 ± 663<br>(877-2678) |
| CLs number (mean±SD) | 0.3 ± 0.5<br>(0-1) | 16.8 ± 6.4<br>(10-29) |
| CTh (mm; range) | 2.72 ± 0.41<br>(2.45-3.15) | 2.53 ± 0.34<br>(2.31-2.99) |
| T2WMLV (cm<sup>3</sup>; range) | 4.7±1.6<br>(1.4-7.2) | 5.6±1.6<br>(3.1-8.4) |

Abbreviations: EDSS: Expanded Disability Status Scale; GM: grey matter; CL: cortical lesion; CTh: cortical thickness; T2WMLV: T2 white matter lesion volume. **Low** and **High** refer to the numbers of cortical lesions on MRI.

