## Supplementary figures and images for "Lymphotoxin-alpha expression in the meninges causes lymphoid tissue formation and neurodegeneration"

### Supplementary Figure 1

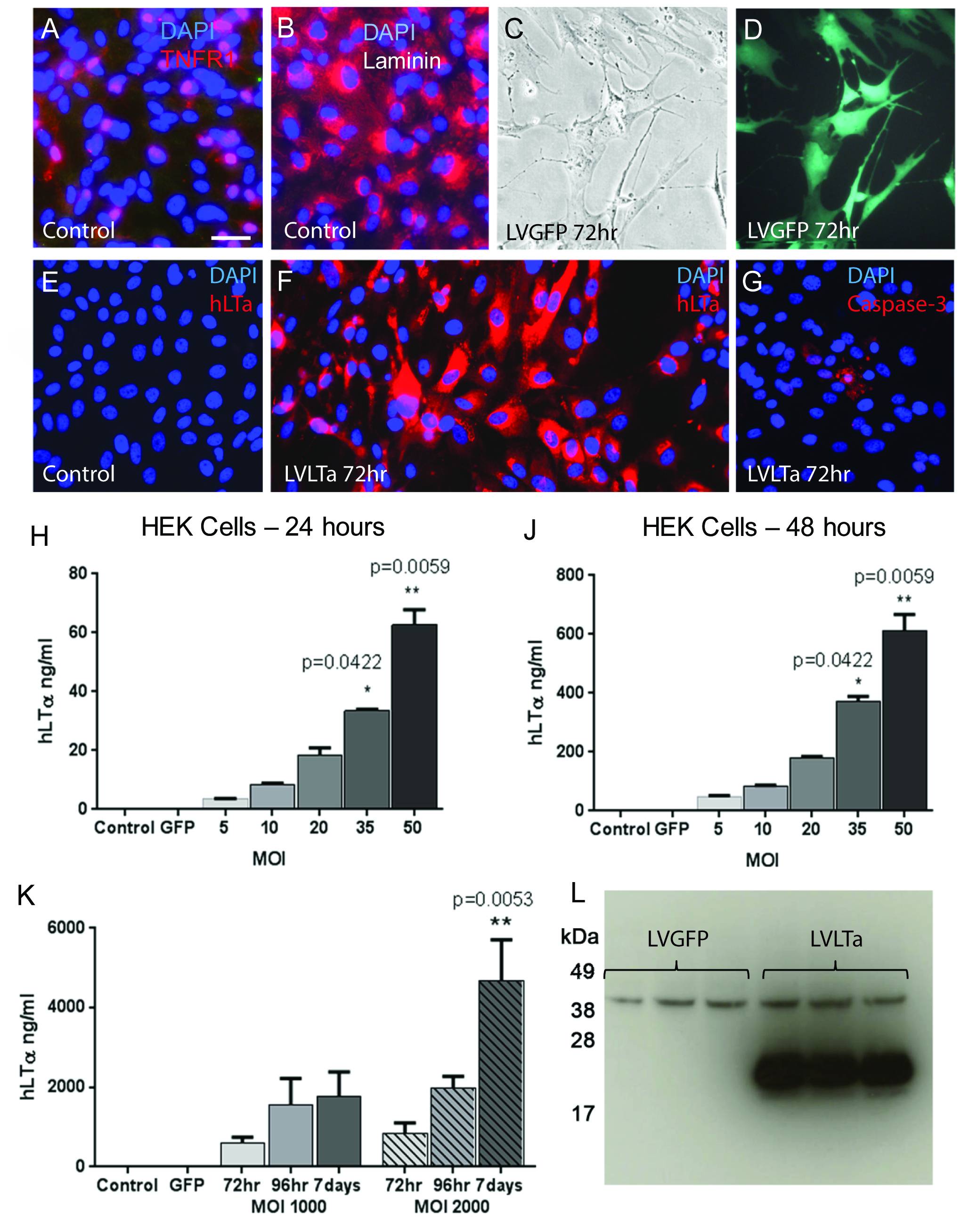
